# Metabolic constraints modulate the likelihood and predictability of global epistasis

**DOI:** 10.1101/2025.02.04.636564

**Authors:** Pavithra Venkataraman, Supreet Saini

## Abstract

Epistasis, where a mutation’s effect depends on genetic background, is central to evolution. A striking pattern, global epistasis, results in fitness effects varying systematically with background fitness—often making beneficial mutations less effective in fitter genotypes. Despite its ubiquity and importance in predicting evolution, the cellular origin of this pattern is unclear. We present a microbial cell growth model and show that global epistasis arises naturally from the underlying metabolic structure, and is modulated by reaction speeds and flux redistribution in the cell. Mutations in standalone rate-limiting modules yield predictable fitness effects, while those in interconnected modules behave idiosyncratically. As genetic divergence increases, background fitness becomes a poorer predictor of fitness effects. Our framework links cellular metabolism to adaptation, offering mechanistic insights into evolutionary predictability.

---

Predicting evolution is a fundamental challenge in biology, with implications on biodiversity, genetics, human health, biogeochemical processes, and agriculture(*1–6*). A key obstacle to predicting evolution is posed by epistasis, whereby fitness effects of mutations in coding and non-coding regions depend on genetic background, and as result, shape evolutionary trajectories in complex ways(*7, 8*). One striking statistical pattern observed in several experimental evolution studies is global epistasis, where the fitness effects of mutations systematically decline as background fitness increases(*9–22*). While this trend allows us to predict epistatic effects of a mutation, there are also exceptions(*14, 23, 24*). The mechanistic origins of these patterns, and when and how they occur, remain unclear(*13, 25, 26*).

Our understanding of global epistasis comes from statistical modeling approaches(*26–28*), or empirically characterized fitness effects of mutations across backgrounds (DFEs)(*12, 17, 29–37*). However, the vast number of possible mutations and genotypes makes full experimental characterization of function and fitness impractical(*38, 39*), and statistical models do not explain how biology structures mutational effects. Metabolic networks and protein interactions have been implicated in epistasis(*40–48*), but no model grounded in cell physiology explains why fitness effects may exhibit global, consistent patterns.

Here, we present a mechanistic framework demonstrating that epistasis and its manifestations arise naturally from the cell’s metabolic structure, allowing us to predict when the fitness effect of a mutation will exhibit global epistasis. Using a quantitative model of microbial metabolism(*49*), we show that the fitness effects of mutations in rate-limiting and standalone enzymatic steps are highly predictable because they are globally epistatic, while mutations in interconnected or upstream modules produce fitness effects that are not predictable using background fitness. We also demonstrate that the extent and nature of genetic background variation strongly influence the globality of epistasis: as background divergence increases, the predictability of a mutation’s fitness effect based on background fitness declines.

These patterns are shaped by the relative speeds of reactions, flux redistribution, and network structure, and persist across metabolic architectures with multiple thresholds and feedback regulation. Our study moves beyond statistical approximations by building a simple yet effective cell growth model grounded in physiology, and explains when and why global epistasis arises. The results generate experimentally testable predictions, and offer a theoretical foundation for understanding how physiology shapes evolutionary constraints.

## Mutational fitness effects and global epistasis

We modelled a prokaryotic cell with three rate-limiting metabolic pathways, each containing two enzymatic reactions (**Fig. 1A, fig. S1-S3, table S1**). Cell division occurs once predefined threshold amounts of all three metabolites (P_1_, P_2_, and P_3_) are accumulated. The time taken to reach this threshold (*t*_*div*_) determines division time, with fitness defined as its reciprocal (1/*t*_*div*_).

**Figure 1.**
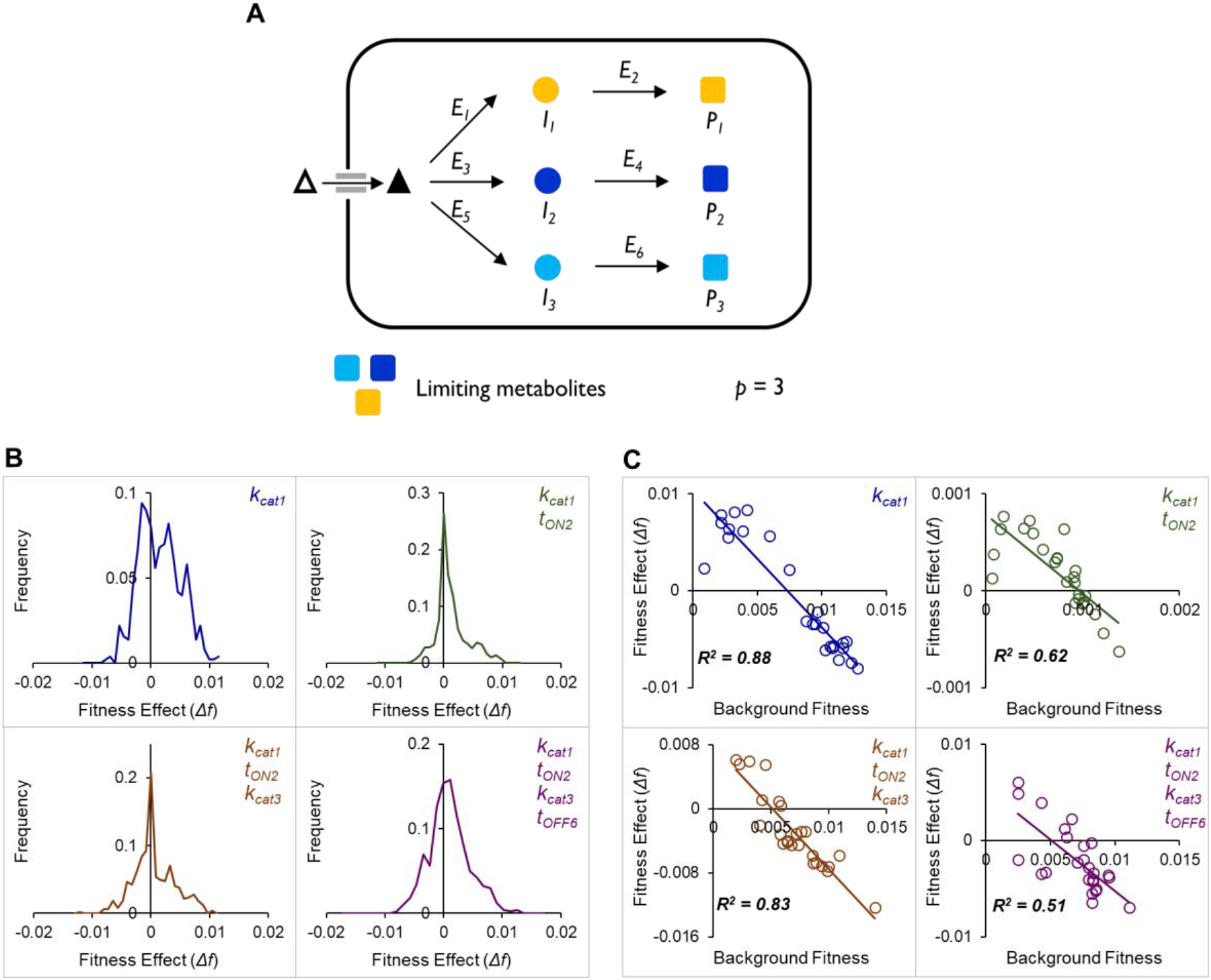
Mutational fitness effects depend on genetic background and vary in predictability. **(A)** Cells divide upon accumulating threshold amounts of three bottleneck metabolites (P_1_, P_2_, and P_3_), each produced via enzymatic cascades from a common substrate. **(B)** Fitness effects of mutations are non-uniform, confirming their dependence on genetic background. The distribution of fitness effects (based on 500 (= 25 × 200) data points) for a fixed mutation (0.5 > *t*_*ON1*_ > 1) is shown across genetic backgrounds differing in one (*k*_*cat1*_), two (*k*_*cat1*_, *t*_*ON2*_), three (*k*_*cat1*_, *t*_*ON2*_, *k*_*cat3*_), or four (*k*_*cat1*_, *t*_*ON2*_, *k*_*cat3*_, *t*_*OFF6*_) background parameters. **(C)** The fitness effect of a mutation is predictable to varying degrees based on background fitness. Regression plots of fitness effects against background fitness for mutations occurring in backgrounds differing by one, two, three, or four parameters.

We call each reaction step a ‘module’, numbered from one to six in the cell. Each module is defined by four parameters. Two of these parameters, *t*_*ON*_ and *t*_*OFF*_, control the promoter activity; while the other two, *k*_*cat*_ and *K*_*M*_, are the enzyme properties (**fig. S2**).

To study epistasis in the context of network-wide interactions, we introduced a mutation by altering the parameter *t*_*ON1*_ (in Module 1) from 0.5 to 1 (represented as 0.5 > *t*_*ON1*_ > 1). This module not only controls the production of intermediate I_1_ but also influences flux through two other reaction pathways by drawing from a shared substrate pool (indicated by a triangle in **Fig. 1A**). This allows us to probe how a single, extreme (the range of *t*_*ON*_ in this work is [0.5, 1]) mutation in an interconnected module interacts with varying genetic backgrounds to produce diverse fitness outcomes, revealing the systemic nature of epistasis. To examine the fitness effect of this mutation across different genetic backgrounds, we introduced it into groups of cells that differ from one another in carefully defined, yet random ways.

Specifically, background variation between any two cells in a group can occur at four positions, relative to the site of the mutation (**fig. S4**): (i) within the same module, (ii) in a different module along the same pathway, (iii) in a module from a different pathway that shares the same substrate, or (iv) in a module from a different pathway that draws from a different substrate. By systematically combining these positions, we arrive at fifteen distinct types of background variation corresponding to all non-empty combinations of one to four varied modules. These variations affect fitness by altering the flux of threshold metabolites. To assess how the extent and nature of background variation shape epistatic interactions and their globality, we generated random variants (analogous to strains) for each of these fifteen types. Within each group, cells differ in a predefined number of parameters. We then introduced the same mutation (0.5 > *t*_*ON1*_ > 1) into each cell and measured its fitness effect. This design allowed us to study mutation effects in backgrounds with unknown internal fluxes. Later in the study (**fig. S5**), we examine mutations in cells with predefined metabolic speeds.

In **Fig. 1B**, we illustrate representative distributions of fitness effects of the *t*_*ON1*_ mutation in four of the fifteen types of background variation types. Distributions for all background variation types are shown in **fig. S6**. For every type of background variation, a distribution was generated by measuring the fitness effect of introducing the *t*_*ON1*_ mutation in twenty-five different cells of twenty independently generated strains. These results reveal that the fitness effects of the same mutation vary substantially across both the type and extent of background variation, indicating that both lower and higher order genetic interactions cause non-additive fitness effects due to changes in the fluxes of the threshold metabolites. This pattern remained robust even when the number of threshold metabolites was altered (**fig. S7**).

A key signature of global epistasis is that the fitness effect of a mutation declines predictably as background fitness increases. To test whether the mutation 0.5 > *t*_*ON1*_ > 1 in Module 1 exhibited this trend, we plotted fitness effects of this mutation against the background fitness. The *R*^*2*^ of this linear regression indicates the globality of epistasis(*50*). Experimental studies have observed epistasis of varying globality in microbial evolution, with *R*^*2*^ ranging from 0.4 to 0.9(*9, 10, 13, 50, 51*). **Fig. 1C** presents representative regression analyses, confirming that our model captures global epistasis. A full set of 300 regressions across all background conditions is provided in **fig. S8**. These regressions have been analyzed in detail in the next section.

## Variability of globality of epistasis

The shape of global epistasis can be characterized by the slope of the regression between fitness effects and background fitness. A negative slope indicates diminishing-returns and increasing-costs epistasis, while a positive slope reflects increasing-returns and diminishing-costs epistasis(*50*). To determine whether global epistasis was predominantly negative, positive, or independent of background fitness, we analyzed the slopes of regression lines. As shown in **Fig. 2A**, negative global epistasis (NGE) remained the dominant pattern across all levels of background variation (**fig. S9, table S2** for statistical comparisons of means). The extent of background variation, however, did not dictate the absolute globality of a mutation, and the mean *R*^*2*^ values remained largely statistically indifferent (**Fig. 2B**, 2-tailed t-test for data with unequal variance was used to compare means; *p >0*.*05* for all but one of the ^4^C_2_ comparisons of means; *p=0*.*023* for the comparison of the mean *R*^*2*^ of 1-site and 4-site variations; see **fig. S10** for full distributions across fifteen background variation types, and **table S3** for statistical comparisons of means).

**Figure 2.**
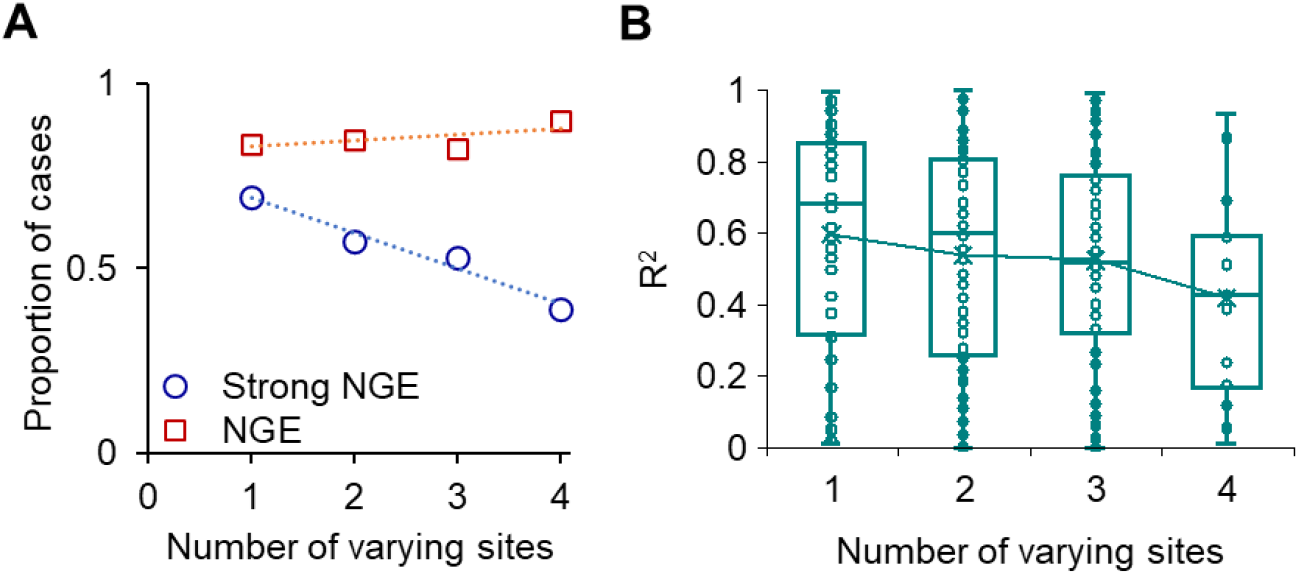
Strong negative global epistasis (NGE) becomes less frequent as background variation increases. **(A)** Negative global epistasis (NGE; squares) remained dominant across all levels of background variation. However, the frequency of strong NGE (*R*^*2*^ > 0.5; circles) decreased as background variation increased. **(B)** The distribution of *R*^*2*^ values for negative global epistasis for different extents of background variation is shown here. Most of the means of the distribution (connected by a line) remained statistically identical (2-tailed t-test for data with unequal variance was used to compare means; *p >0*.*05* for all but one of the 6 comparisons of means; *p=0*.*023* for the comparison of the mean *R*^*2*^ of 1-site and 4-site variations). See **Figure S10** for full distributions across all 15 background variation types.

We defined negative, strongly global epistasis (strong NGE) as cases where *R*^*2*^ > 0.5, distinguishing them from weaker instances. Using this threshold, we quantified how the globality of epistasis changed with changes in the levels of background variation. As shown in **Fig. 2A**, the likelihood that a mutation exhibits strong NGE decreases, as the genetic divergence among the cells in which it is introduced, increases.

## Metabolic basis of the globality of epistasis

We next asked whether the globality of epistatic effects of a mutation can be predicted from the relative speeds of enzymatic modules within a metabolic network. If certain reactions are slow (rate-limiting), they constrain overall pathway flux, and mutations in these steps could have more predictable effects across different backgrounds. In contrast, when metabolic speeds vary widely or multiple steps compete for shared substrates, the effects of a mutation can become highly context-dependent and idiosyncratic, potentially weakening the predictability implied by global epistasis. Thus, metabolic speeds shape how genetic background modulates the impact of mutations.

To understand if metabolic speeds dictate the globality of epistasis, we constructed four types of simplified (“toy”) cells where six enzymatic modules were predefined as either fast or slow (see Methods 1.5). As shown in **fig. S5**, we created the following variants:

- 1A – First-step module is slow; all others are equally fast.
- 1B – First-step module is fast; all others are equally slow.
- 2A – Second-step module is slow; all others are equally fast.
- 2B – Second-step module is fast; all others are equally slow.

Thus, each cell type contains one outlier reaction step compared to the rest. We simulated twenty-five cells of each type and introduced a hundred random mutations in the outlier step to assess how global epistasis operates.

Only mutations in the slow standalone module (2A) exhibited strongly global epistasis, with fitness effects tightly correlated with background fitness (**Fig. 3A**). Mutations in other configurations (1A, 1B, and 2B) displayed weak global epistasis patterns. We also computed the slope and intercept of the regression of fitness effect v/s background fitness for each mutation, and examined their relationship with the *R*^*2*^ of the fit (a measure of globality). In case 2A, slope and intercept were strongly correlated with *R*^*2*^ (r = 0.85 and –0.93, respectively), indicating that the strength and magnitude of mutational effects are tightly coupled to the degree of globality (**Fig. 3B–3E**). The only other strong correlation (|r| > 0.8) was observed in case 2B between intercept and *R*^*2*^ (r = – 0.84), though *R*^*2*^ values were generally low in this case (*R*^*2*^ < 0.5).

**Figure 3.**
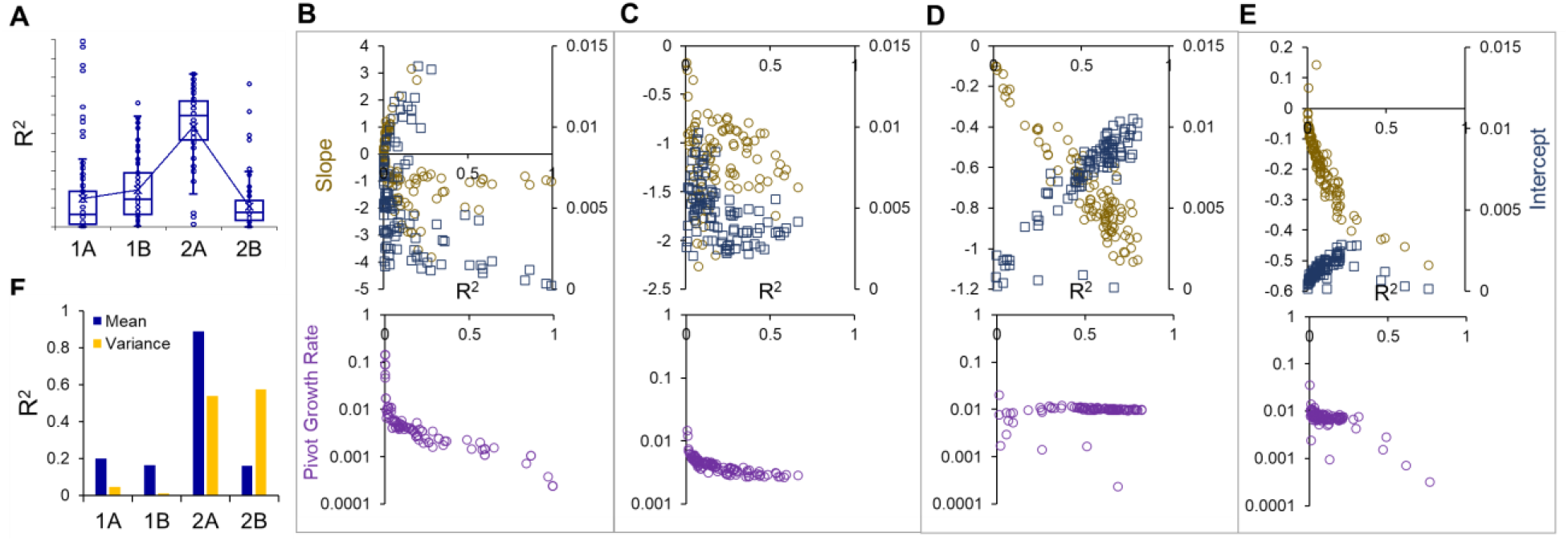
Predictability of mutational fitness effects depends on reaction speed and metabolic control. **(A)** Mutations in 2A exhibit more global epistatic effects than mutations in the other modules. Shown here are the *R*^*2*^ values of a linear fit between the fitness effects of the mutations and the background fitness for a hundred mutations in twenty-five different genetic backgrounds of the four cell types (1A, 1B, 2A and 2B) (See **fig. S5** for the design of the four cell types). **(B-E)** Slope and intercept are strongly coupled with the globality in case 2A (r = 0.85 and –0.93, respectively). For each of the hundred regressions in the four cell types, we measure the slope (circles, brown), intercept (squares, blue), and pivot growth rate (circles, purple) (cases 1A, 1B, 2A, and 2B are shown in **(B), (C), (D)**, and **(E)**, respectively). When epistasis is strongly global (*R*^*2*^ *>* 0.5), pivot growth rate is predictable in the slow modules – it decreases with an increase in globality in case 1A, and remains invariant in case 2A. **(F)** In case 2A, the mean of the DFE, and variance of the DFE in cases 2A and 2B are well predicted by the background fitness. In each genetic background, we computed the mean and variance of the DFE of the hundred mutations. The mean and variance of the DFE of a background was obtained by considering the fitness effects of the hundred mutations. The *R*^*2*^ values of these linear regressions are shown.

We next examined the pivot growth rate—the background fitness at which a mutation’s effect changes sign—a quantity determined by the slope and intercept of the regression. In strongly global cases (*R*^*2*^ > 0.5), pivot growth rates were highly predictable for mutations in the slow modules. In case 1A, pivot growth rate decreased with increasing globality (r = –0.87), whereas in case 2A, it remained invariant (r = 0.00086). In cases 1B and 2B, the rarity of strongly global epistasis precluded meaningful inference. These findings suggest that pivot growth rate is invariant only for mutations in slow and standalone reaction steps, supporting recent claims that pivot behavior may be environmentally robust(*13*).

Finally, we analyzed the distribution of fitness effects (DFEs) for each of the twenty-five genotypes. The mean of the DFE consistently declined with increasing background fitness in all four cell types (**fig. S11**), but this relationship was most predictable in case 2A (**Fig. 3F**). The variance of the DFE increased with background fitness in cases 2A and 2B (**Fig. 3F, fig. S12**), but not in 1A or 1B, suggesting that mutations in standalone reaction modules may enhance evolvability at high fitness(*52*).

Mutations in the slow and standalone reaction module continued to exhibit global epistasis when the metabolic network was reduced two threshold metabolites (each synthesized via a three-reaction cascade) (**fig. S13**), and also when strong feedback inhibition was added (i.e., pathways shut down upon reaching metabolite thresholds; **fig. S14-S15**). These results confirm that mutations in rate-limiting standalone modules exhibit global, and hence, predictable fitness effects.

Together, these findings offer a mechanistic explanation for how metabolic structure constrains or enables global epistasis, and this framework helps explain empirical observations that beneficial mutations become rarer at high fitness, while deleterious effects become more prevalent(*13, 17, 53*)—an outcome now grounded in pathway dynamics.

## Metabolic modularity and fitness effects

Mutations in fast, non-rate-limiting modules are typically expected to have weak or nearly neutral fitness effects because they do not constrain pathway flux. As a result, such mutations are non-epistatic and do not exhibit global epistasis. However, in slow reaction modules, mutations are expected to have consistent and predictable fitness effects because they strongly constrain pathway output and, consequently, fitness. However, our results show that mutations in 1A do not yield predictable fitness effects.

In a branched or interconnected network, a mutation in the first slow step (1A) can alter flux distribution across multiple pathways. This redistribution introduces context-dependence and may reduce the predictability of fitness effects by breaking the clean global epistasis pattern observed in standalone bottlenecks. We tested this by analyzing the effect of four predefined mutations (in *k*_*cat*_) introduced into six variants each of 1A- and 2A-type toy cells (see **Methods, Section S1.5**). For each case, we measured whether the fluxes of three threshold metabolites (P_1_, P_2_, P_3_) changed significantly due to the same mutation, under otherwise identical parameters.

As shown below in **Fig. 4A**, due to a mutation in *k*_*cat2*_ (case 2A), rates of production of P_2_ and P_3_ significantly changed only in 6.25% of the instances, while mutations in *k*_*cat1*_ (case 1A) caused a change in rates of P_2_ and P_3_ in 25% of the instances, indicating greater flux redistribution. We also measured the rate of change of fitness per rate of change of *k*_*cat*_ (sensitivity, *f*_*k*_) and examined how it varies with background fitness, to identify if mutations in 1A were epistatic at all. As shown in **Fig. 4B**, *f*_*k*_ and fitness show a strong negative correlation, for both cases 1A and 2A (Refer to **fig. S16** for the regressions). However, the slope of this correlation is much more negative in the case 2A, compared to case 1A (**Fig. 4C**, *p < 0*.*001*, 1-tailed t-test for unequal variance), indicating a stronger global component of fitness effects of mutations in 2A.

**Figure 4.**
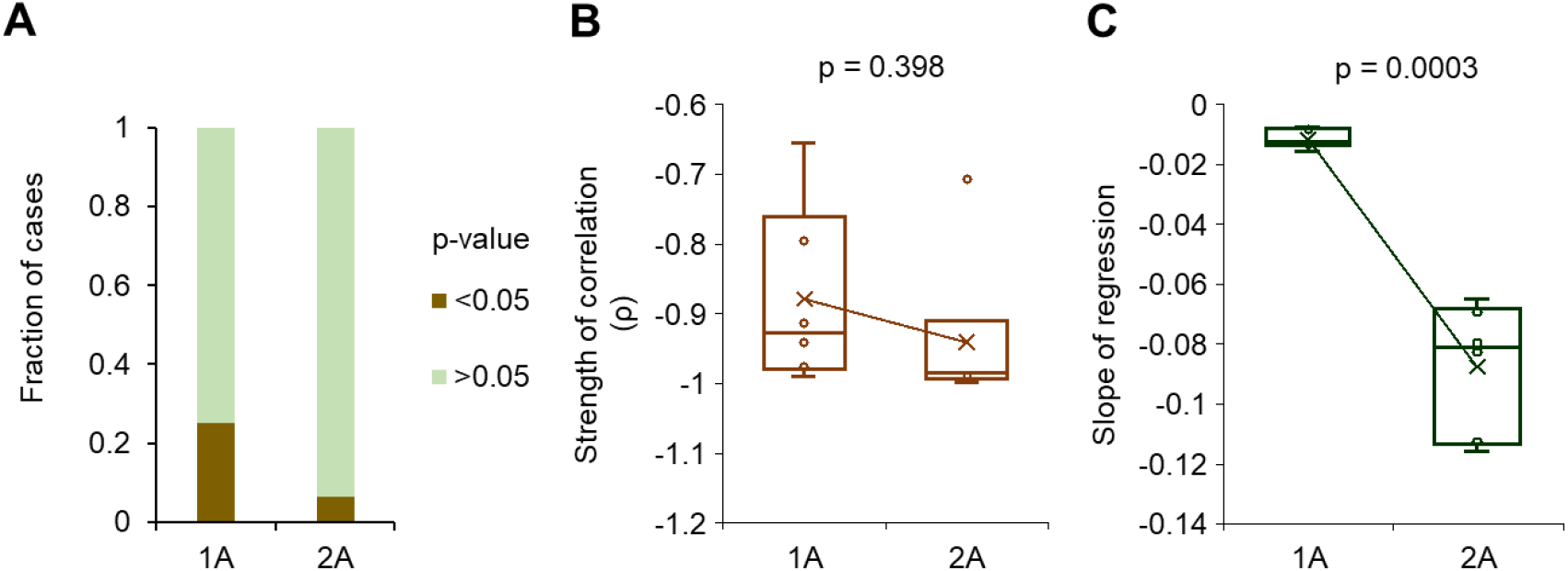
The effects of mutations in interconnected reaction modules are spread across pathways. **(A)** Mutations in 1A lead to a significant change in the rates of production of P_2_ and/or P_3_ four times more frequently than mutations in 2A. We measured the effect of four large effect mutations (in *k*_*cat*_), in six backgrounds each, on the rate of production of the three threshold metabolites, in 1A and 2A types of cells. This indicates that the fitness effects of mutations in 1A are more likely to affect all three pathways. **(B)** We next asked how the fitness of the cell changes due to a unit change in the mutation parameter (*k*_*cat*_), by measuring the ratio of the change in fitness to the change in *k*_*cat*_ (*f*_*k*_). *f*_*k*_ and background fitness are strongly negatively correlated in both cases 1A and 2A, as shown by the Pearson’s r in the figure. The *p-value* was obtained by comparing the means of the average r in the two cases using a 2-tailed t-test for unequal variance. **(C)**. The slope of the linear regression is highly non-negative and variable for mutations in 2A, compared to the mutations in 1A, as shown in **Figure S16**. The *p-value* reported in this figure was obtained by comparing the means of the average slopes in the two cases using a 2-tailed t-test for unequal variance.

Despite the decline in sensitivity of fitness to mutations with increasing background fitness, global epistasis does not manifest strongly in 1A where mutations cause flux redistribution across multiple pathways (**Fig. 4A**). This results in the possibility of similar fitness values despite different internal flux states and genotypes. Consequently, mutation effects vary among backgrounds with comparable fitness, weakening the correlation between mutation effect and background fitness, reducing predictability using global epistasis. However, in 2A, flux redistribution is minimal. Hence, similar fitness corresponds to similar internal metabolic states, leading to more consistent mutation effects and stronger, more predictable global epistasis. Thus, although mutations in 1A influence fitness, their effects are diluted and modulated by network interactions, explaining the weaker global epistatic fitness effects for these mutations. These results highlight how metabolic network architecture influences the predictability of evolutionary trajectories by shaping the relationship between genotype, flux distribution, and fitness.

## Discussion

In this study, we develop a mechanistic model of cell growth and division that captures epistasis and its globality, revealing how metabolic constraints shape mutational effects and their predictability. Fitness in our model is defined by the time required to accumulate threshold levels of key metabolites before cell division. Cellular metabolic networks are often highly interconnected, with many reaction steps often influencing flux through other pathways—except for terminal steps that lead directly to metabolite production(*54*). While our model simplifies this complexity, it includes two distinct types of reaction modules: those that influence flux through other branches, and those that do not. This design captures the essential dichotomy present in real metabolic networks—interconnected vs. standalone modules—and allows us to study epistasis in a comprehensive and mechanistically grounded manner, including the effects of both coding and non-coding mutations.

Empirical studies show that negative global epistasis (NGE)—where the effects of beneficial mutations diminish as background fitness increases—is common but not universal(*14, 25*). Our model recapitulates this trend and predicts that negative epistasis becomes less global as the tested genetic backgrounds become more divergent in the parameter space. We also show that mutations in the slowest, standalone reaction modules consistently produce globally epistatic fitness effects. As these slow steps improve, beneficial mutations become rarer, and are expected to contribute to declining adaptability over time(*17, 34, 53, 55*).

Beyond its theoretical insights, this framework has broad implications for understanding how epistasis operates. Importantly, our model addresses a key limitation of the recent statistical theories of epistasis(*26, 28*), which explain global epistasis as a statistical consequence of many uncorrelated genetic interactions and regression to the mean, without accounting for the physiological mechanisms that underlie fitness. In contrast, we ground genotype–fitness relationships in metabolic physiology by modeling how mutations affect enzyme activities and, in turn, steady-state metabolic fluxes—a biologically meaningful intermediate phenotype. In this structured, non-random interaction framework, global epistasis emerges from non-linear flux– fitness mappings, such as saturation effects or pathway bottlenecks, rather than from statistical averaging alone. By incorporating metabolic constraints, our model provides a mechanistic basis for global epistasis that is both predictive and interpretable in terms of underlying cellular processes. Future work could extend this model by including cellular regulation layers and environmental changes, to explore the role of these factors in dictating epistasis’ predictability(*50, 56*). Currently, our findings could help in engineering microbial strains and communities with predictable evolutionary trajectories, impacting biotechnology and antimicrobial resistance research(*32, 57, 58*).

Our model generates clear testable predictions. In the metabolic map of the cell, several key metabolites feed branches of enzymatic reactions. For instance, when *E. coli* is grown in anaerobic conditions, acetyl-CoA feeds into the Krebs cycle, fatty acid biosynthesis, and mixed acid fermentation(*59–61*). Similarly, glutamate is a substrate for the synthesis of glutamine, proline and arginine(*62*). Our model predicts the reaction modules in these pathways in which mutations, under appropriate conditions, would exhibit strong global epistasis. Our work also predicts sign-epistasis between promoter and gene mutations, which is likely to constrain evolutionary trajectories through mechanisms which are not fully understood yet (**fig. S3A, S17**)(*56, 63*).

Ultimately, this work provides a quantitative foundation for predicting fitness gains in evolving populations. Future studies will be crucial in testing and refining these insights, to further our understanding of the role of cellular processes in dictating evolutionary dynamics.

## Supporting information

Contains Methods and Supplement Figures.

## Acknowledgments

We are grateful to Deepa Agashe, Sunil Laxman, Sandeep Krishna, Christian Landry, Sergey Kryazhimskiy, Narendra Dixit, and Kirti Jain for their insights and feedback. We also thank Disha Chaudhary and Sarvesh Baheti for their comments on the manuscript.

## Author contributions

Conceptualization: PV, SS

Methodology: PV, SS

Investigation: PV, SS

Funding acquisition: PV, SS

Project administration: SS

Supervision: SS

Writing – original draft: PV, SS

Writing – review & editing: PV, SS

## Data and materials availability

All data are available in the main text or the supplementary materials. Raw data of all figures and Matlab codes used in this study are available at: https://github.com/SainiSupreet/Global-Epistasis

