## Supplementary material for "Metabolic constraints modulate the likelihood and predictability of global epistasis": Contains Methods and Supplement Figures.

### 1. Methods

#### 1.1 Cell growth and division model based on a metabolic adder and sizer

We model cell growth and division based on the production of three bottleneck metabolites. These bottleneck metabolites are produced via independent two-step enzymatic reactions, all of which are fed by a substrate drawn from the environment, as shown in **Figure S1**. When a cell accumulates predefined amounts ($10^8$ molecules in this work) of all the threshold metabolites, it divides, and each of the daughter cells inherits half of all the cellular constituents (mRNA, enzymes, and metabolites). The time taken for a cell to divide is $t_{div}$, and $\frac{1}{t_{div}}$, the growth rate, is considered a proxy for the fitness of the cell.

It was shown previously that the model captures the division time statistics observed in an isogenic population of a microbial species(*49*). The model is based on a metabolic sizer/adder principle of microbial cell division, which is known to statistically capture division times and sizes in bacteria and yeast(*64*, *65*).

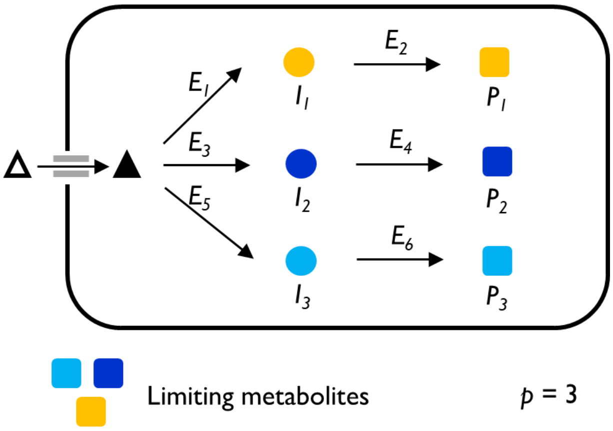


**Figure S1. Essential metabolic pathways in the cell.** The cells modelled in this work divide when they accumulate threshold amounts of limiting metabolites. A cell with three bottleneck metabolites ($P_1$, $P_2$, and $P_3$) is shown. Each terminal metabolite is produced via enzymatic cascades, feeding on the same substrate.

#### 1.2 Stochastic gene expression and production of enzymes

Independent gene-promoter systems dictate the production of the enzymes that drive the reactions in the cell, as shown in **Figure S2**. The process of gene expression comprises of stochastic (burst-like) transcription, and deterministic translation. To implement stochastic-burst-like gene expression, we sample random "$t_{ON}$" and "$t_{OFF}$" durations of each of the promoters from two exponential distributions of set means. In this study, the means of $t_{ON}$ and $t_{OFF}$ are drawn from uniform distributions $U_1(0.5,1)$ and $U_2(0.5,5)$, respectively(*49*, *66*). The process of transcription proceeds in a deterministic fashion for $t_{ON}$ duration. Each mRNA is assigned a lifetime duration drawn from an exponential distribution of a set mean (= 25 minutes)(*67*). Only one ribosome is attached to an mRNA at a given instant of time to translate it. We sample an exponential distribution of a set mean (= 20 hours) to assign lifetime durations for each protein molecule produced(*68*). The average length of the gene is 1029 bp(*69*), the rate of transcription is 10 nucleotides/s, and the rate of translation is 10 amino acids/s(*70–72*).

Every enzyme involved in the production of the intermediate and threshold metabolites is characterized by a $k_{cat}$ and $K_M$, values of which are drawn from uniform distributions $U_3(1,100)$ and $U_4(6022, 602200)$ respectively, based on data reported in the literature(*73*).

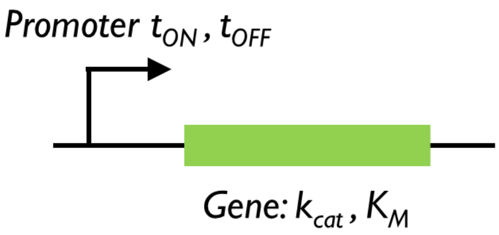


**Figure S2. Gene and promoter characteristics.** Every enzyme involved in the production of threshold metabolites is produced based on the characteristics of its promoter. We model stochastic, burst-like transcription. The enzyme catalysis is dictated by $k_{cat}$ and $K_M$.

##### 1.3 Production of metabolites via enzymatic reactions

Michaelis-Menten reaction laws govern the conversion of one metabolite to another. The substrate S, denoted by Δ in **Figure S1**, is common for the three reaction arms. The reactions occurring in the cell are modelled according to the differential equations below.

$$\frac{dS}{dt} = 30000 - \frac{k_{cat1}E_1S}{K_{M1}+S} - \frac{k_{cat3}E_3S}{K_{M3}+S} - \frac{k_{cat5}E_5S}{K_{M5}+S} \quad (1)$$

$$\frac{dI_1}{dt} = \frac{k_{cat1}E_1S}{K_{M1}+S} - \frac{k_{cat2}E_2I_1}{K_{M2}+I_1} \quad (2)$$

$$\frac{dP_1}{dt} = \frac{k_{cat2}E_2I_1}{K_{M2}+I_1} \quad (3)$$

$$\frac{dI_2}{dt} = \frac{k_{cat3}E_3S}{K_{M3}+S} - \frac{k_{cat4}E_4I_2}{K_{M4}+I_2} \quad (4)$$

$$\frac{dP_2}{dt} = \frac{k_{cat4}E_4I_2}{K_{M4}+I_2} \quad (5)$$

$$\frac{dI_3}{dt} = \frac{k_{cat5}E_5S}{K_{M5}+S} - \frac{k_{cat6}E_6I_3}{K_{M6}+I_3} \quad (6)$$

$$\frac{dP_3}{dt} = \frac{k_{cat6}E_6I_3}{K_{M6}+I_3} \quad (7)$$

The amount of each metabolite that the cell must accumulate before it divides is set at $10^8$, capturing division time distributions observed in prokaryotes(*49*).

A simulation of metabolite synthesis and accumulation, via stochastic transcription and deterministic translation and enzyme kinetics is shown in **Figure S3**. In this simulation, cell division takes place upon accumulation of $10^8$ molecules of the terminal metabolite. Upon division, the two daughter cells inherit half the transcripts, protein molecules, and metabolites. Due to stochastic transcription, cell division times exhibit a distribution in an isogenic population[1].

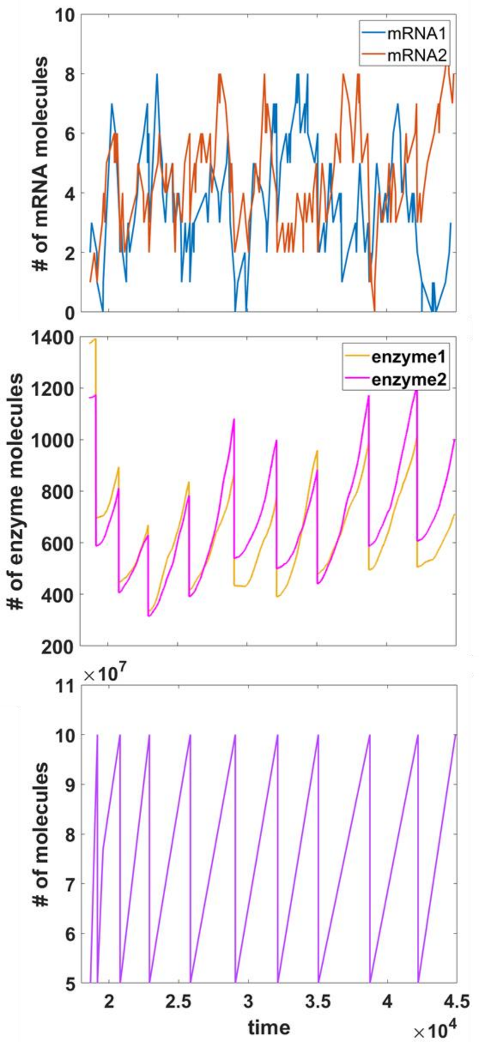


**Figure S3. Temporal profiles of mRNAs, enzymes, and threshold metabolite.** We model a cell with stochastic transcription, deterministic translation, and division based on accumulation of threshold metabolites. In this scenario, for one of the three enzymatic cascades, the temporal profiles of the two mRNAs (top panel), two enzymes (middle panel), and the threshold metabolite (bottom panel) are shown.

##### 1.4 Creation of background variation

As shown above, the cells we model have six enzymes (*3 threshold metabolites x 2 reactions to produce each of them*). Each of these enzymes' amounts and catalytic properties are governed by four parameters – $t_{ON}$, $t_{OFF}$, $k_{cat}$, and $K_M$. Therefore, the fitness of a cell is a non-linear function of twenty-four parameters.

In this section, unless specified otherwise, we study the effect of a mutation in $t_{ON1}$ (0.5 to 1), on fitness. We call this the focal mutation, and study its effect on fitness ($\Delta f = f_{tON1=1} - f_{tON1=0.5}$) in different genetic backgrounds.

To create genetic variations in the cell, one could vary any of the 23 parameters (while $t_{ON1}$ is the 24$^{th}$) which dictate a cell's fitness. However, in a module, varying a gene or a promoter has a similar distribution of fitness effects (**Figure S17**).

Hence, for the focal mutation in a given module (module 1 in our study), we could have background variation via mutations in - (a) the same module, (b) a different module in the same path, (c) a module in the same position but different path, and (d) a module in a different path and position. For the focal mutation in $t_{ON1}$, we create the background variation by changing (a) $k_{cat1}$ (same module), (b) $t_{ON2}$ (different module in same path), (c) $k_{cat3}$ (same position but different path), and (d) $t_{OFF6}$ (different module and position). All background variation is thus generated by changing these one or more of these four parameters only **(Figure S4)**.

Cells where background variation is introduced via changing only one (of $k_{cat1}$, $t_{ON2}$, $k_{cat3}$, or $t_{OFF6}$) parameter are of four types $\binom{4}{1}$. Similarly, cells where background variation is introduced by changing (a) two parameters is $\binom{4}{2}$; (b) three parameters is $\binom{4}{3}$; and (c) all four parameters is $\binom{4}{4}$.

Thus, the different combinations of these classes yield fifteen ($\binom{4}{1} + \binom{4}{2} + \binom{4}{3} + \binom{4}{4}$) types of background variations that could be created.

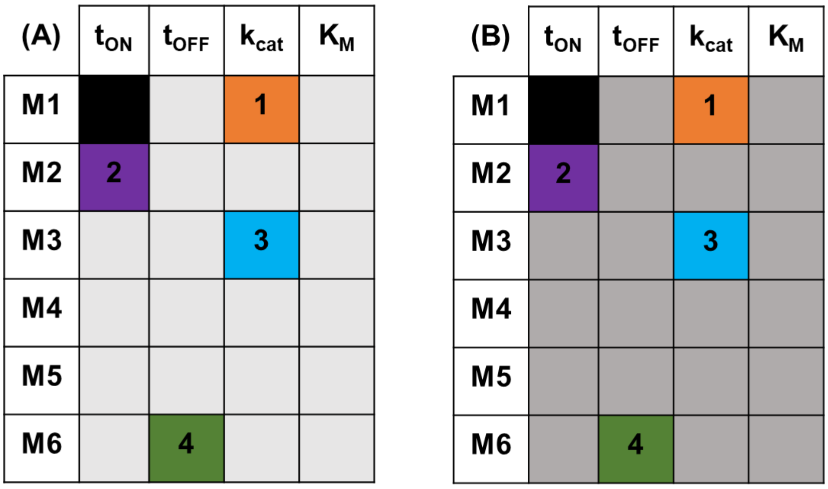


**Figure S4. Variations in the parameters to create different backgrounds.** In this figure, the rows indicate the six modules in the cell, and the columns indicate the four parameters in each of these modules. In this study, a mutation in $t_{ON1}$ (0.5 to 1, cell highlighted in black in **(A)** and **(B)**) has been studied in different "genetic" backgrounds. In order to create these different backgrounds, four parameters are varied in different combinations, while keeping the others constant. The four parameters are $k_{cat1}$ (indicated by 1 in this study, highlighted in orange in **(A)** and **(B)**), $t_{ON2}$ (indicated by 2 in this study, highlighted in violet in **(A)** and **(B)**), $k_{cat3}$ (indicated by 3 in this study, highlighted in yellow in **(A)** and **(B)**), and $t_{OFF6}$ (indicated by 4 in this study, highlighted in green in **(A)** and **(B)**). The effect of variations in the four parameters is studied in different backgrounds, indicated by different shades of grey for the remaining 19 parameters in **(A)** and **(B)**. The different shades of grey indicate changes in the macro interactions, and we equate such changes with differences between strains.

Consider the case where there is one varying parameter, say $t_{ON2}$. In order to study the epistatic effects of lower-order interaction, we create a pool of twenty-five cells which differ in $t_{ON2}$, identical to each other in twenty-two parameters, and the twenty-fourth parameter is the focal mutation. This group of cells can be thought of as those belonging to the same strain, varying at only one site. (Similarly, for cases where there are two, three, and four varying parameters, twenty-one, twenty, and nineteen parameters are kept constant, respectively).

To study the epistatic effects of higher-order interactions, we create twenty such distinct pools (these can be thought of as twenty different strains), and each pool has a unique set of twenty-two parameters. We then introduce the focal mutation and study its fitness effects in these resulting five hundred cells.

We study the effect of the focal mutation using this approach in all the fifteen background variations.

##### 1.5 Creation of cells with predefined metabolic speeds

We create cells with predefined metabolic speeds to study global epistasis. The speed of a reaction module is either "slow", "same" or "fast" relative to the other reactions **(Figure S5)**. In this study, we create toy cells in which the $t_{OFF}$ and $K_M$ of all the six modules are identical and equal to 5 and 60220 respectively. $t_{ON}$ of all six modules are identical to each other, but is a value between 0.5 and 1. This variation in $t_{ON}$ gives cells of different background fitness. A reaction step's speed, relative to the others, is decided by its $k_{cat}$, which is 10 for "slow" reactions, and 100 for "fast" reactions.

In each of these odd module backgrounds, we then introduce a hundred random mutations in $k_{cat}$, and measure their fitness effects ($\Delta f = f_{mutant} - f_{background}$) in twenty-five different backgrounds. We use this dataset to study the likelihood of global epistasis of mutations in these four types of reaction modules.

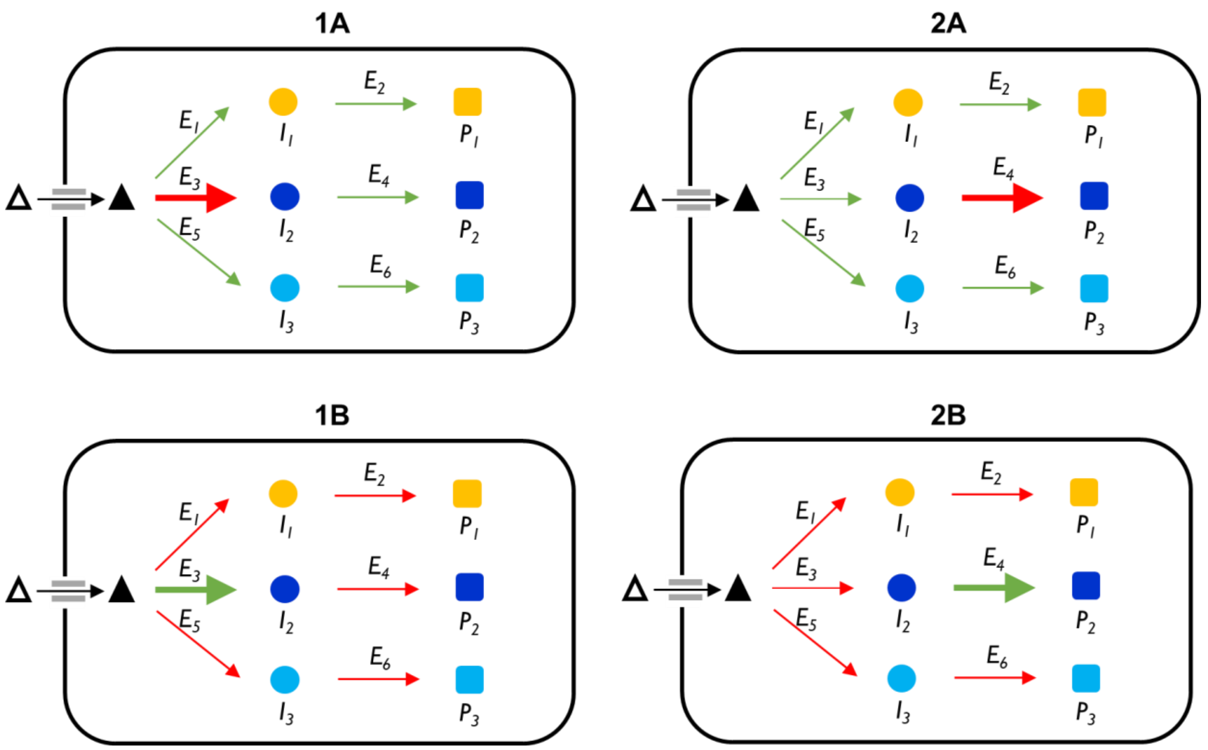


**Figure S5. Toy cells with predefined metabolic speeds.** The cells that we model have three reaction arms which produce the three threshold metabolites. Each of the reaction arms have two reaction steps. In total, there are six reaction modules. We create toy cells in which the speeds of each of these modules is predefined as either slow or fast, relative to the others. Arrows in red and green indicate slow and fast reactions, respectively. In **1A** type of cells, one of the first reaction steps is slow, while the others are all equally fast. In **1B**, one of the first step reactions is fast, and the other five are all equally slow. In **2A**, one of the second step reactions is slow and the other five are equally fast, and in **2B**, one of the second step reactions is fast and the other five modules are equally slow.

To test the effect of mutations in 1A and 2A on the fluxes of $P_1$, $P_2$, and $P_3$, we constructed toy cells with predefined metabolic speeds as described above, but with non-randomly chosen $t_{ON}$ values and mutations. Specifically, we analyzed the fitness effects of four mutations in $k_{cat1}$ (for case 1A) and $k_{cat2}$ (for case 2A), with values changing from 10→20, 20→40, 40→60, and 60→80. These mutations were introduced into six different genetic backgrounds, which differed only in $t_{ON}$ values (0.5, 0.6, 0.7, 0.8, 0.9, and 1). In all backgrounds, the non-mutated modules were fast ($k_{cat}$ = 100), and $t_{OFF}$ and $K_M$ were set to 5 and 60,220 respectively. The $t_{ON}$ values for all operons were uniformly set to one of the six background values.

For each of the 1A and 2A cases, we measured the effects of the four mutations (averaged over 100 cells) on the fluxes of $P_1$, $P_2$, and $P_3$, as well as on overall fitness, across all six backgrounds. The rate of production for each metabolite was calculated as the difference in molecule count between the start and end of the cell cycle, divided by division time. To determine whether a mutation caused a statistically significant change in production rate, we performed a two-tailed t-test assuming unequal variances on the rates before and after mutation. A *p-value* $< 0.05$ was considered indicative of a significant effect.

To calculate $f_k$ values reported in the main text, we divided the change in fitness by the corresponding change in $k_{cat}$ for each mutation. These $f_k$ values were then plotted against background fitness to examine the background-dependence of mutational effects.

##### 1.6 Simulations

All simulations were performed in MATLAB R2022a. All codes used in this study are given at https://github.com/SainiSupreet/Global-Epistasis.

#### 2. Supplementary Figures

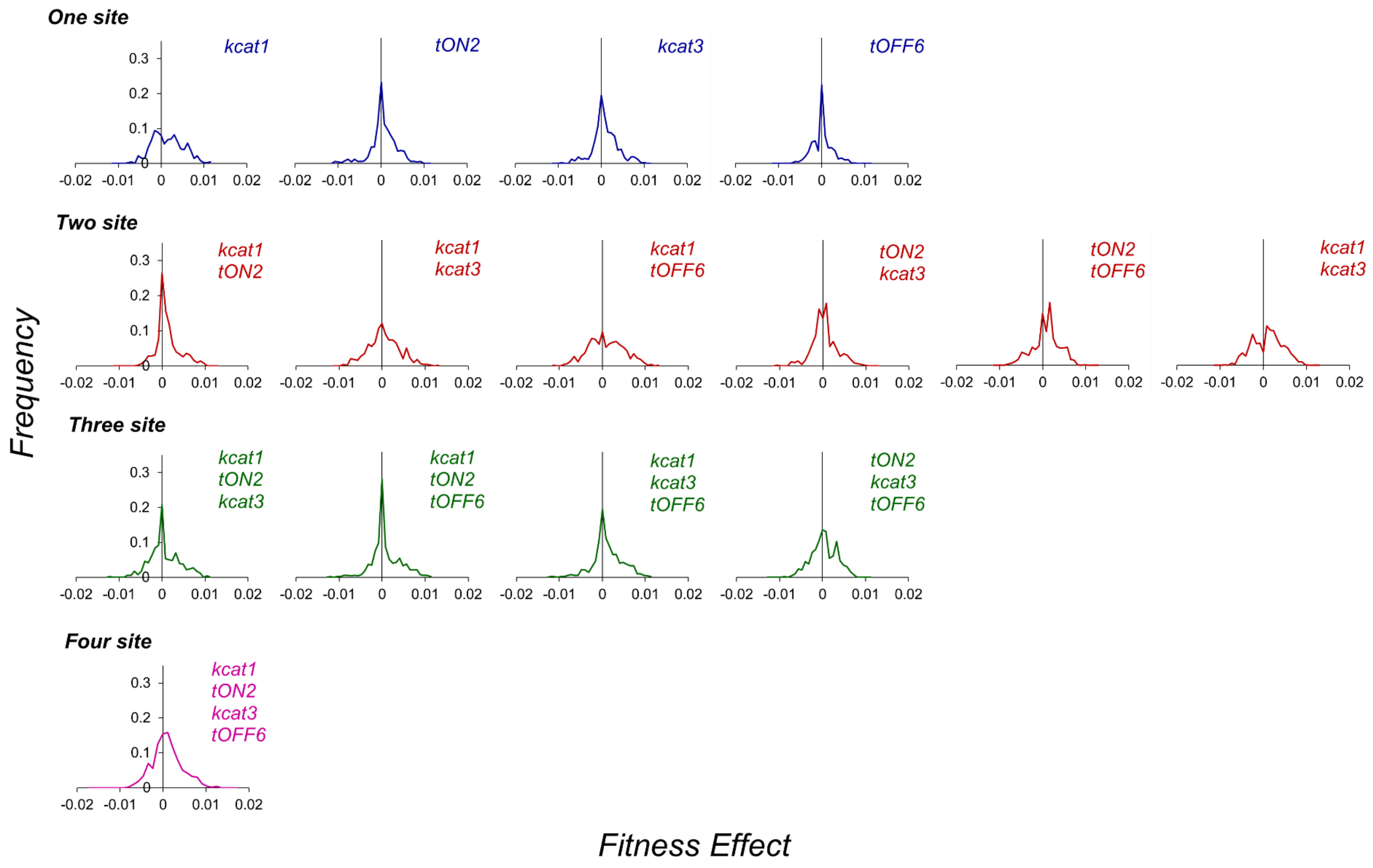


**Figure S6. The fitness effects of mutations are background-dependent.** We draw up the distribution of fitness effects of a fixed mutation ($t_{ON1}$ changes from 0.5 to 1) when it occurs in a group of cells that differ from each other in one, two, three and four background sites.

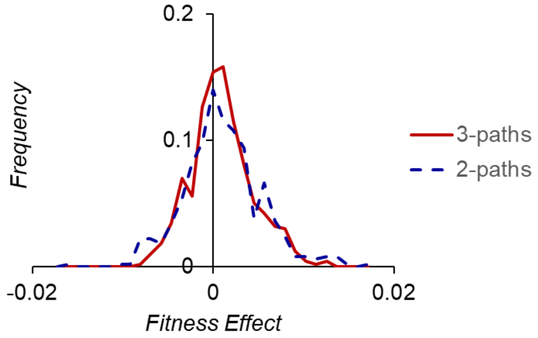


**Figure S7. Changes to the metabolic architecture do not alter background dependence of fitness effects.** The fitness effects of mutations change with change in the genetic background in which they occur, in a cell with three threshold reaction arms, and two reaction steps in each. We confirm that background-dependent effects are not exclusive to this metabolic architecture by studying the effect of a mutation in $t_{ON1}$ in a cell with two threshold reaction arms with two reaction steps in each. As shown in the figure above, fitness effects of mutations remain non-uniform across genetic backgrounds.

A. Variation in $k_{cat1}$

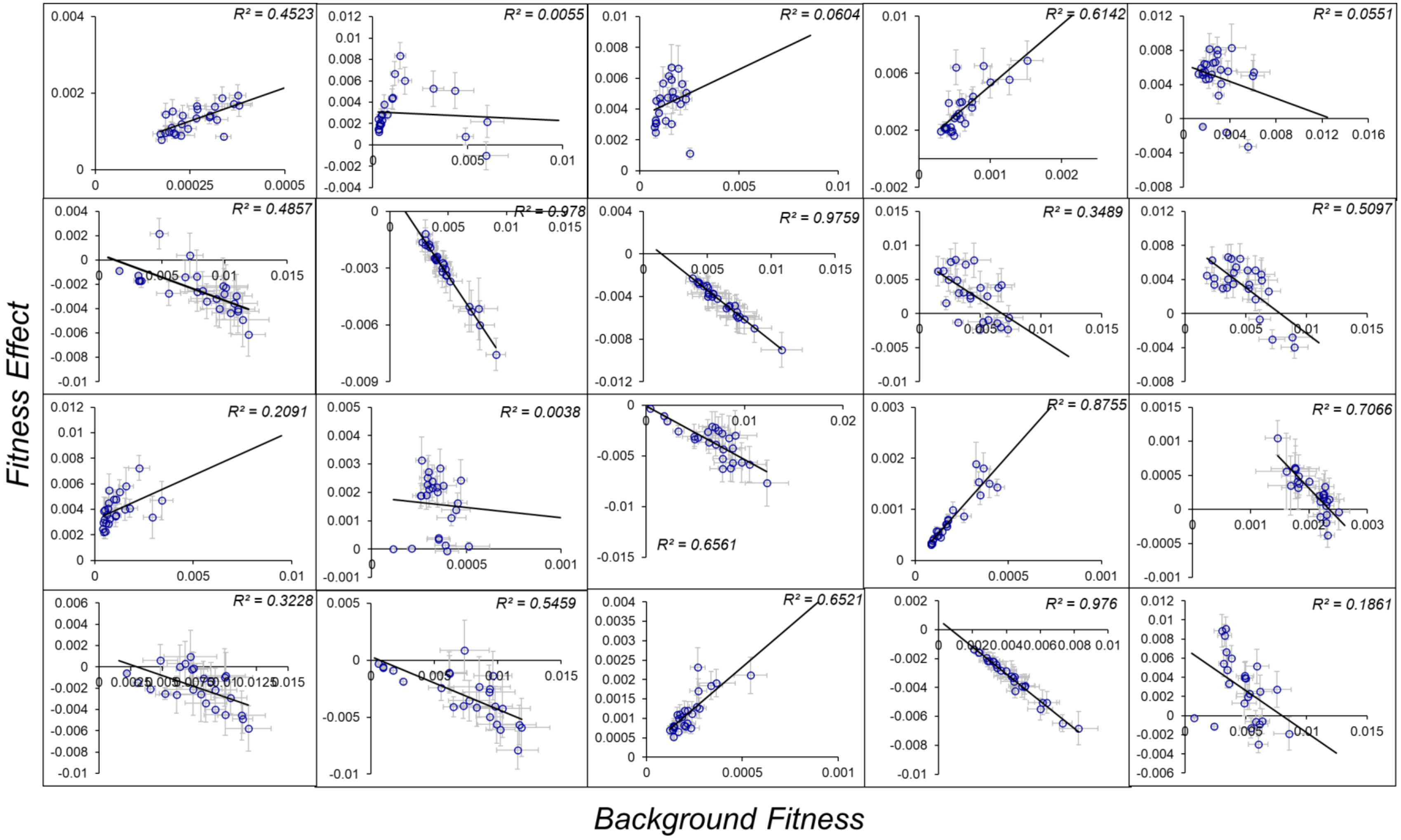

*B.* Variation in $t_{ON2}$

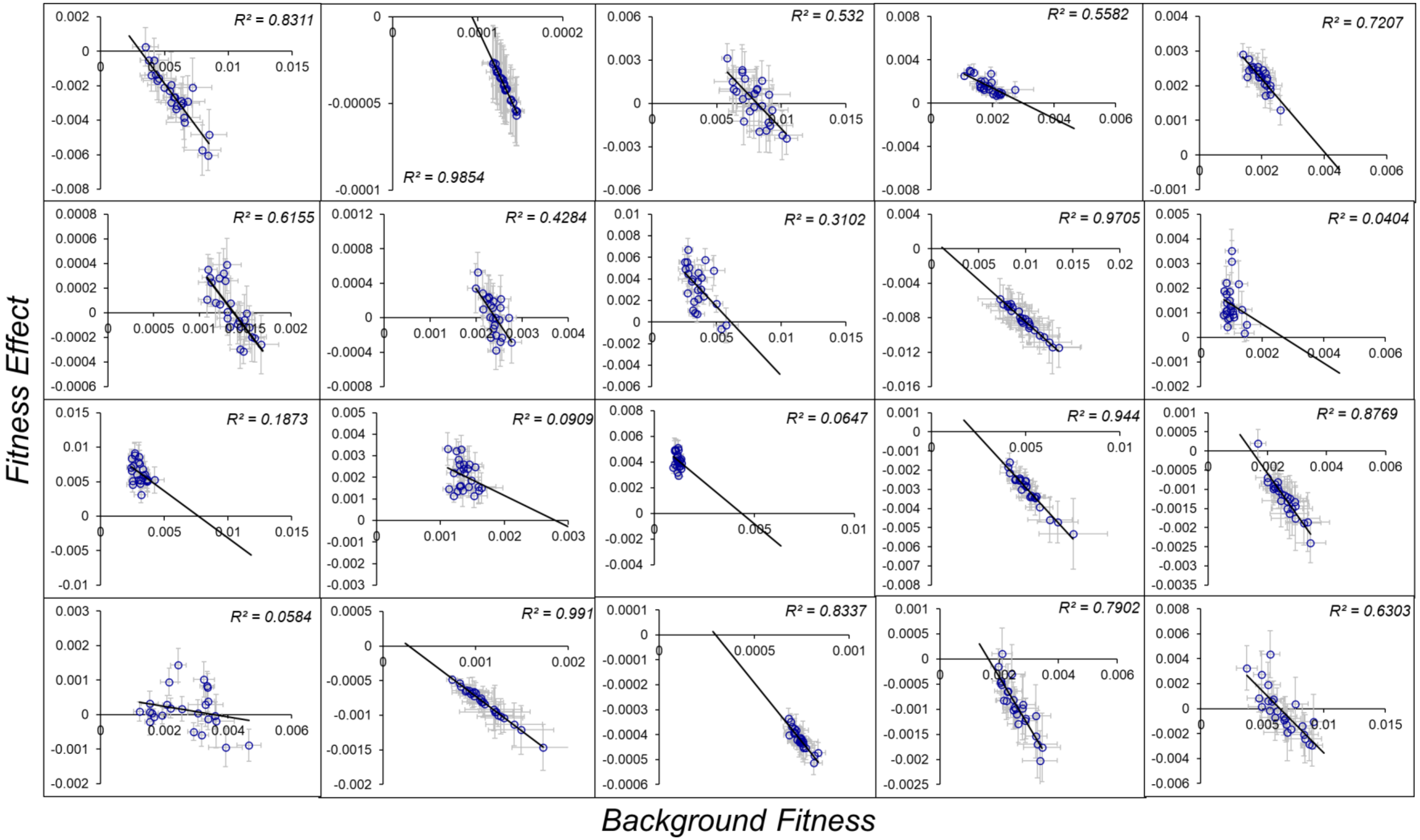

C. Variation in $k_{cat3}$

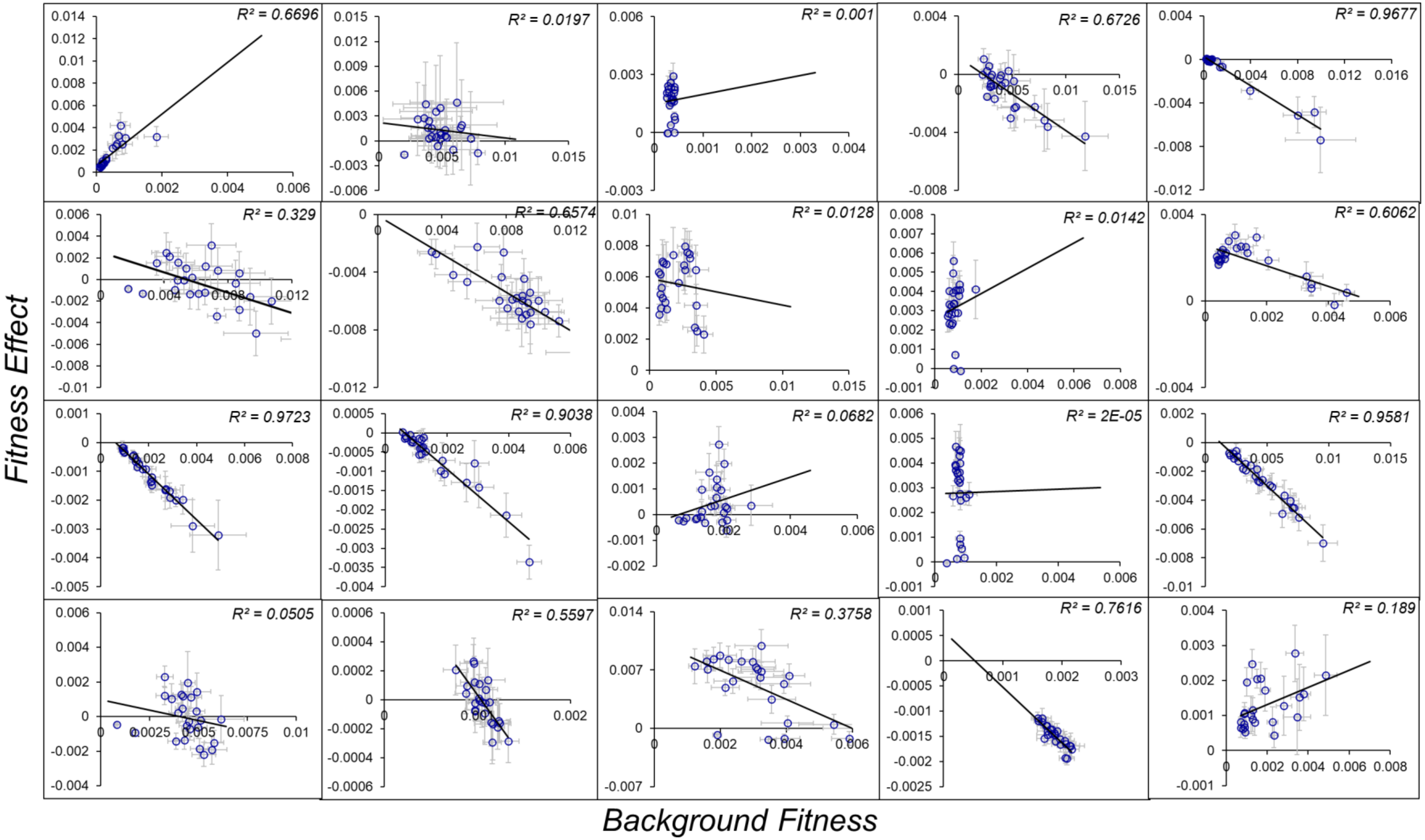

D. Variation in $t_{OFF6}$

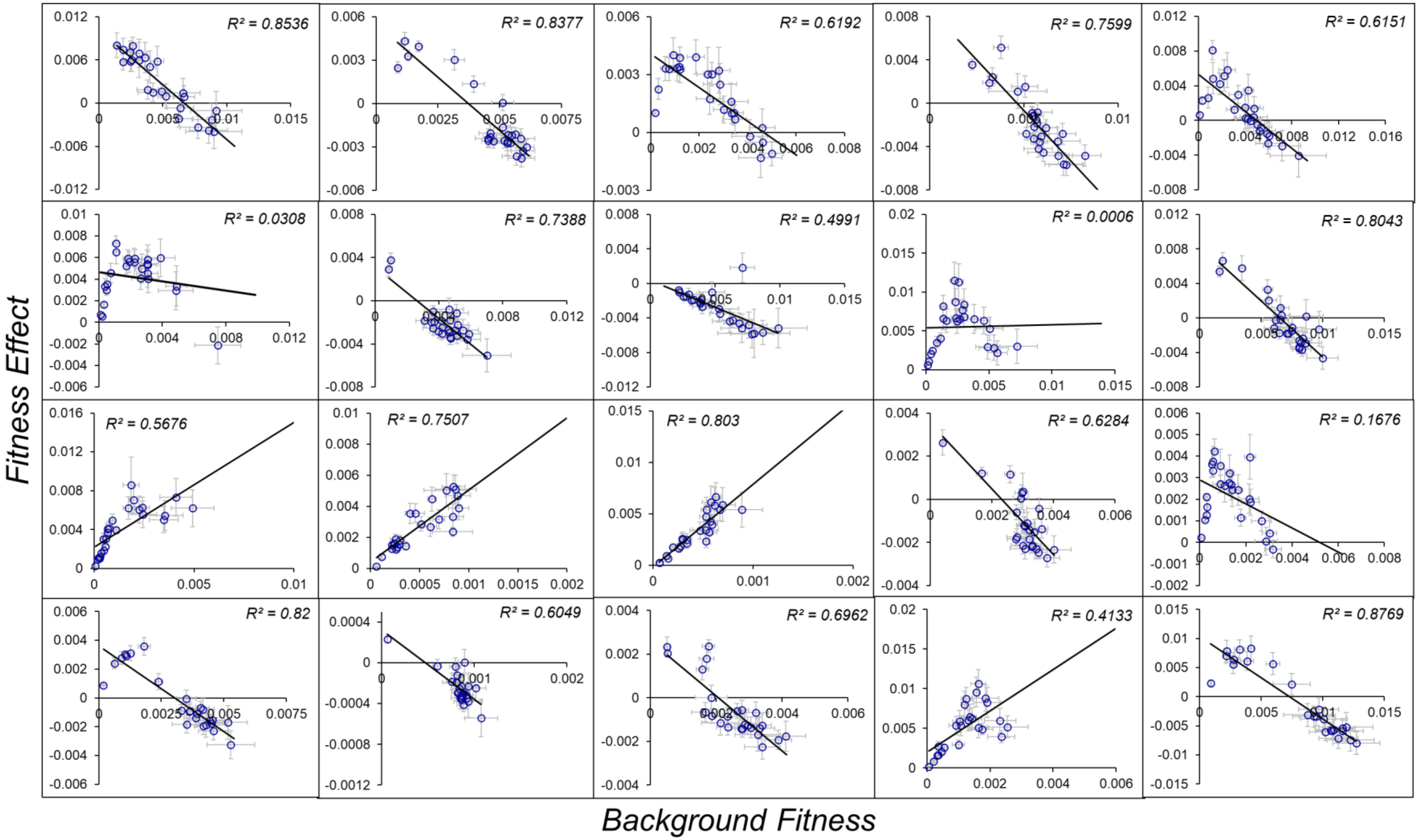

E. Variation in $k_{cat1}$ and $t_{ON2}$

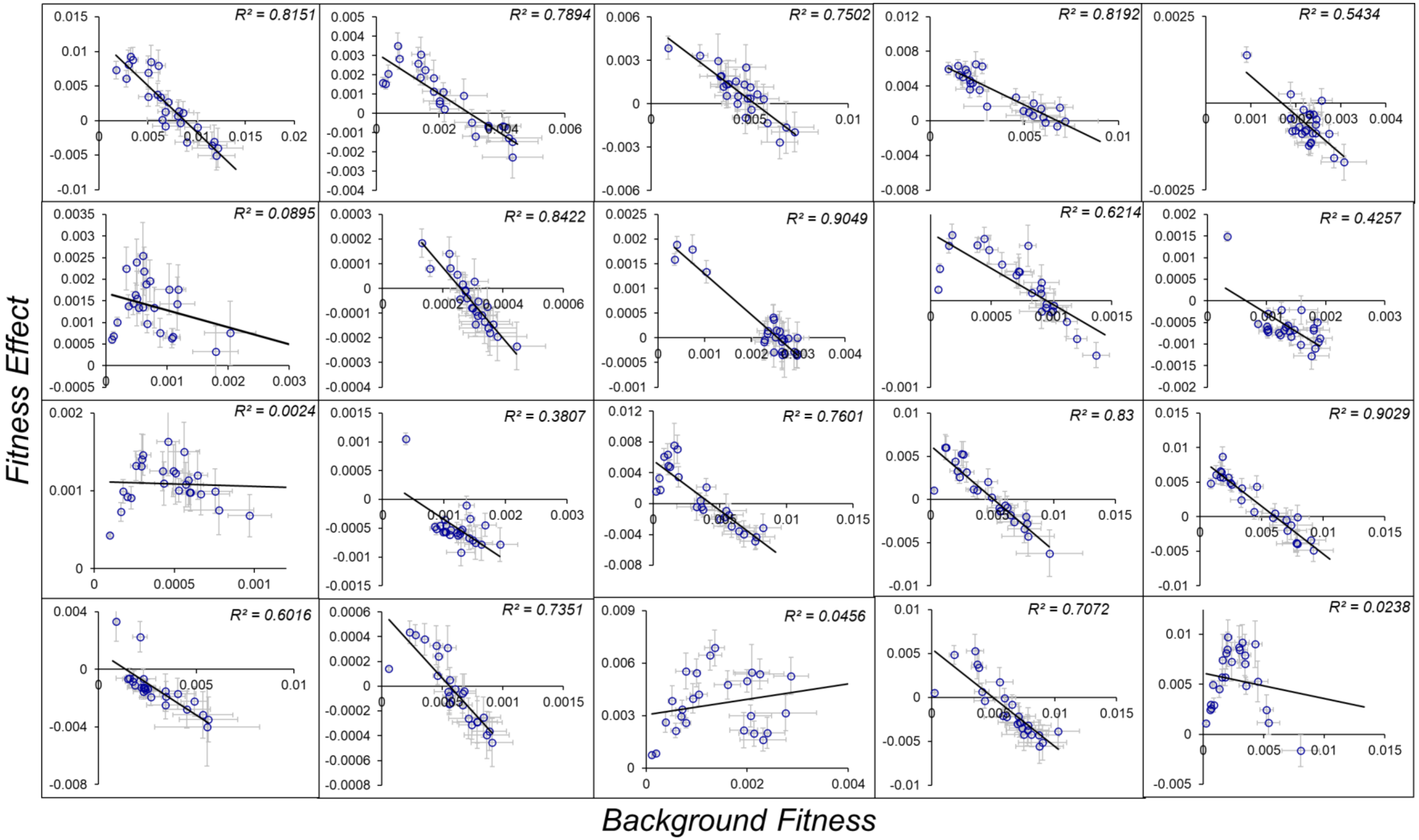

F. Variation in $k_{cat1}$ and $k_{cat3}$

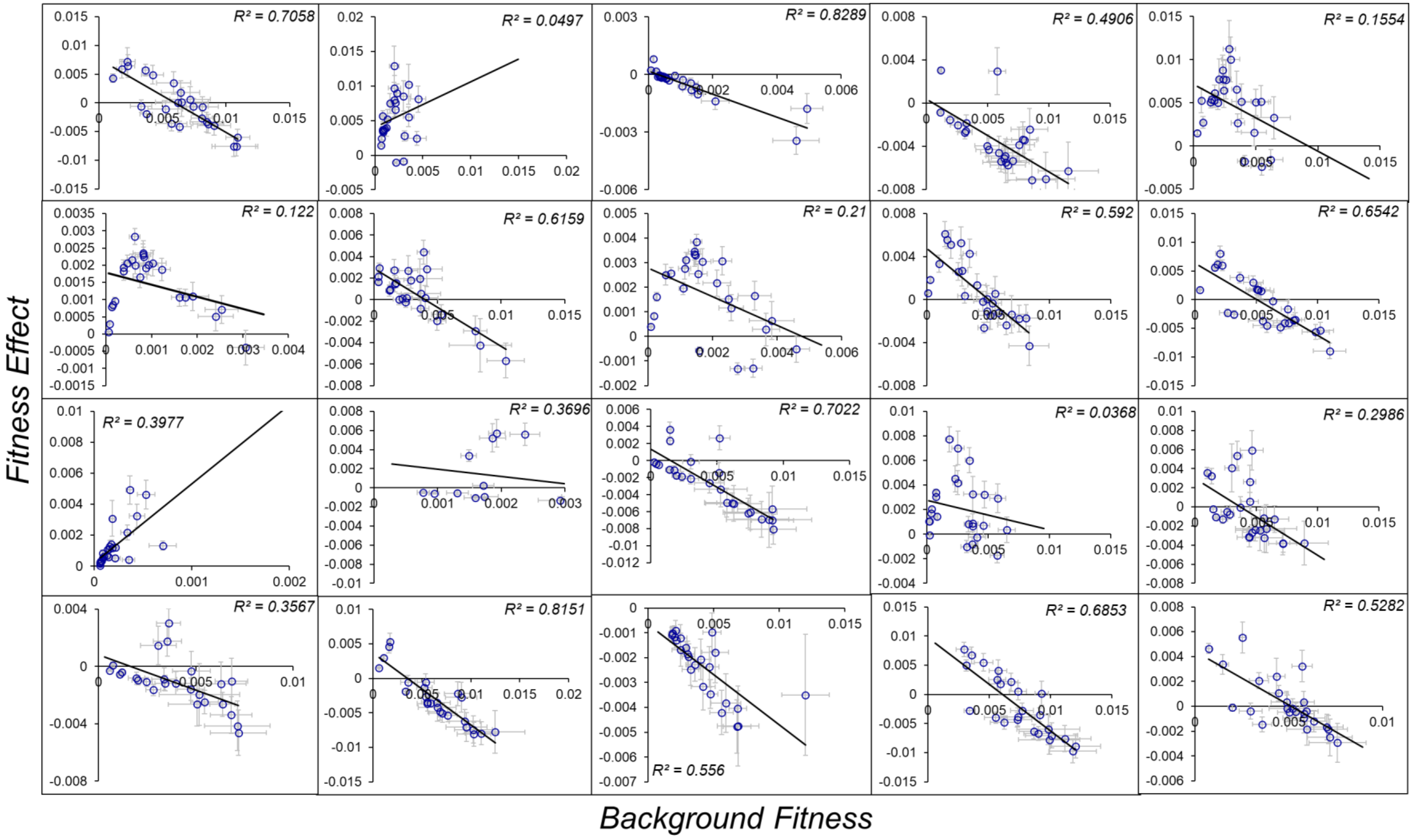

G. Variation in $k_{cat1}$ and $t_{OFF6}$

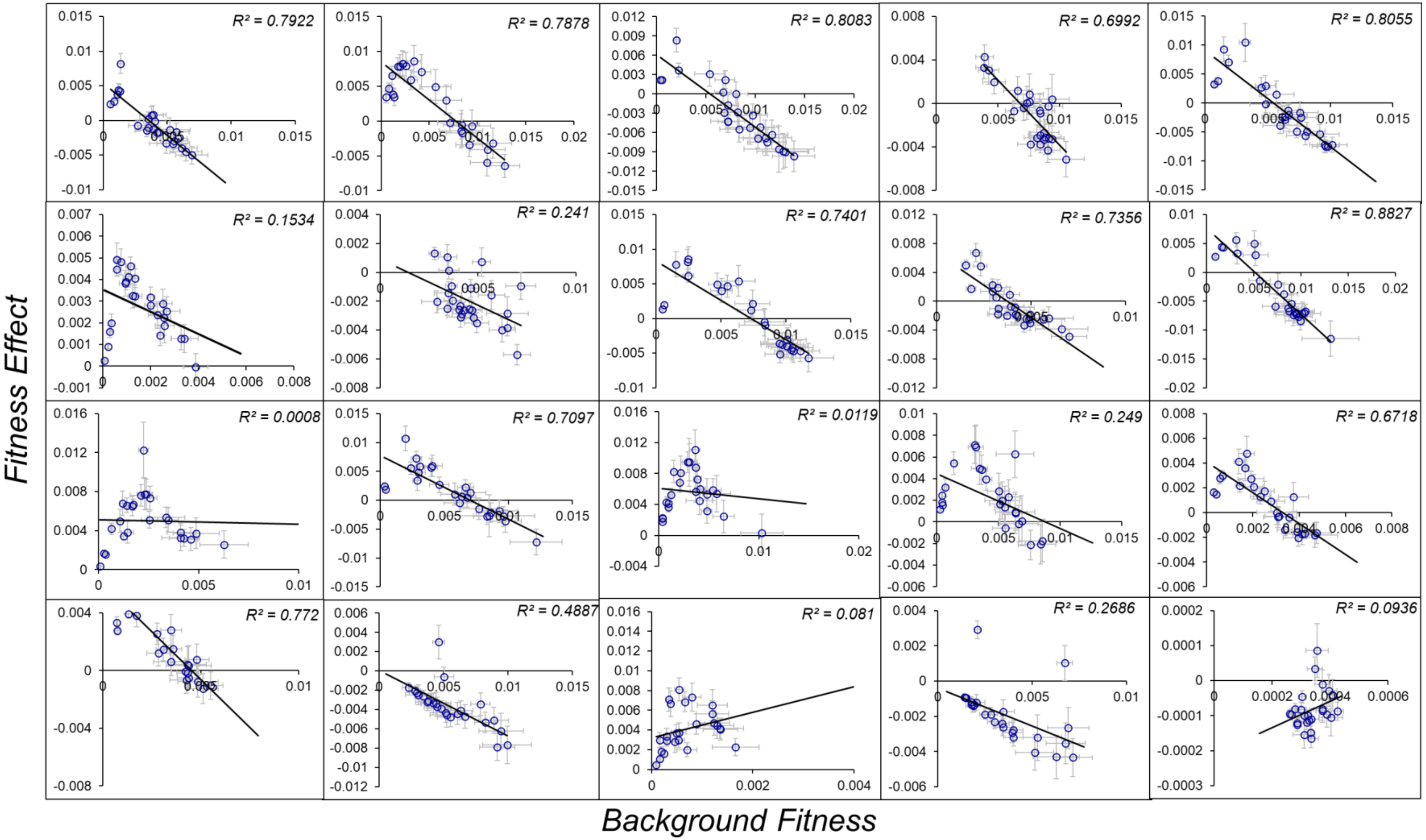

H. Variation in $k_{cat3}$ and $t_{ON2}$

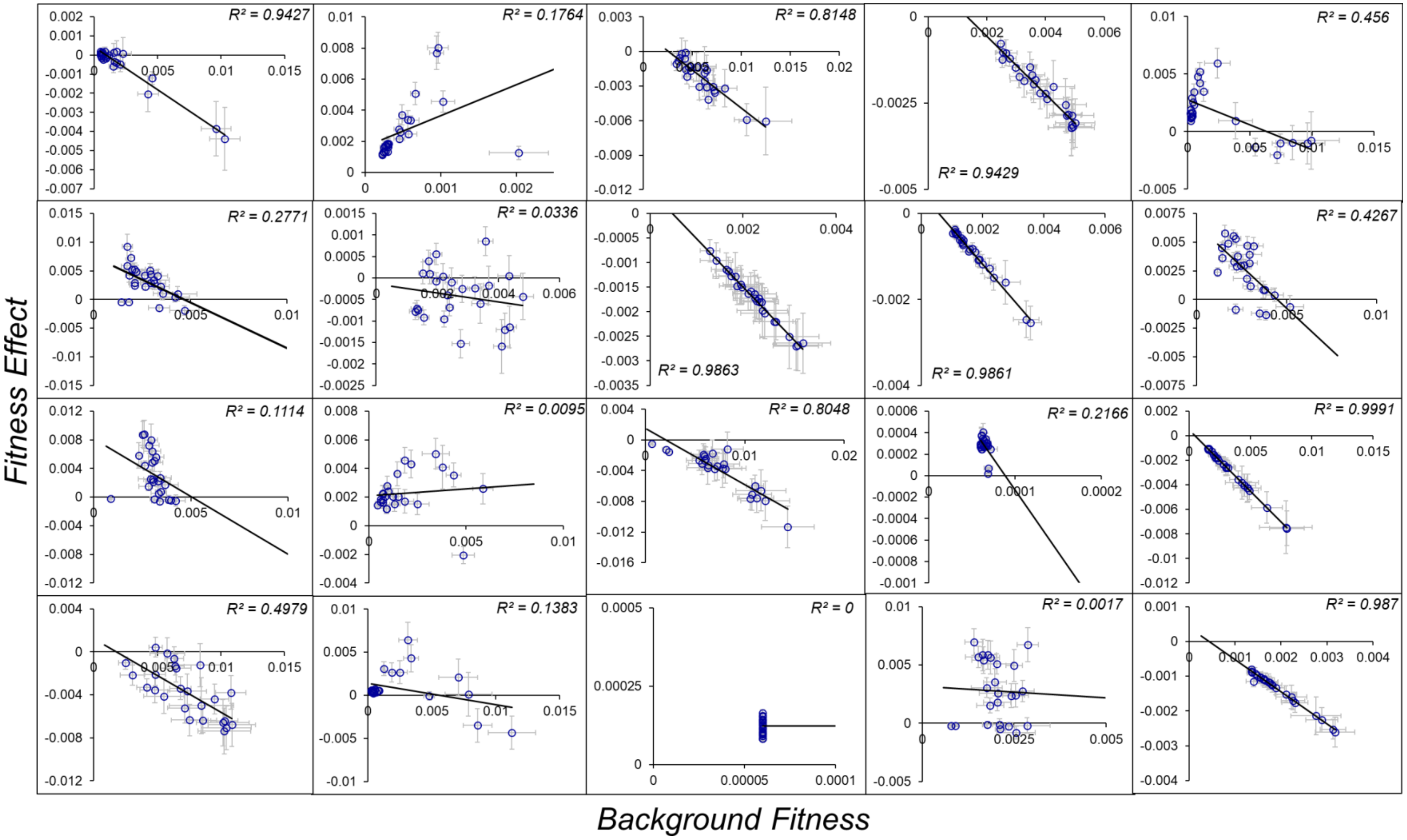

I. Variation in $t_{OFF6}$ and $t_{ON2}$

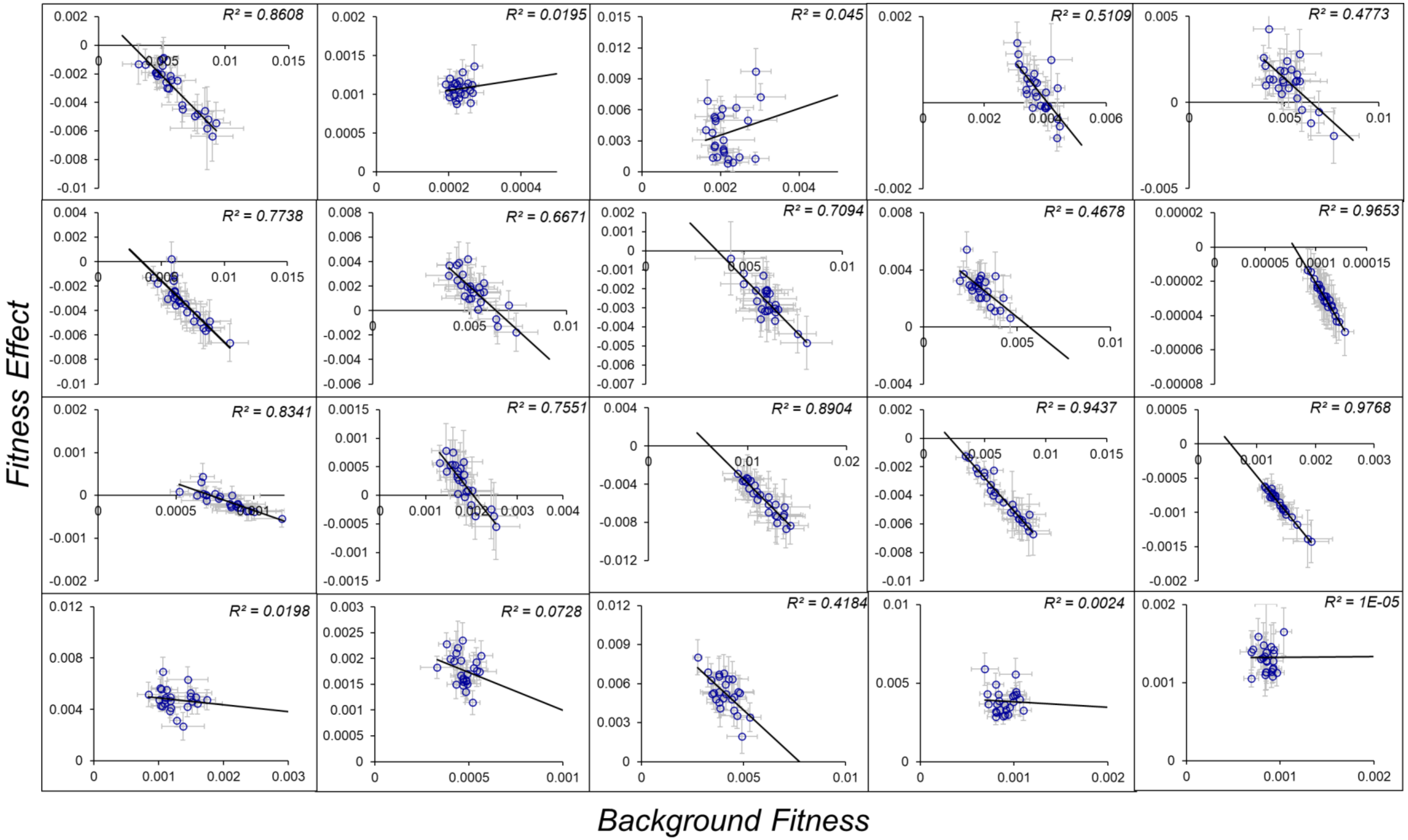

J. Variation in $t_{OFF6}$ and $k_{cat3}$

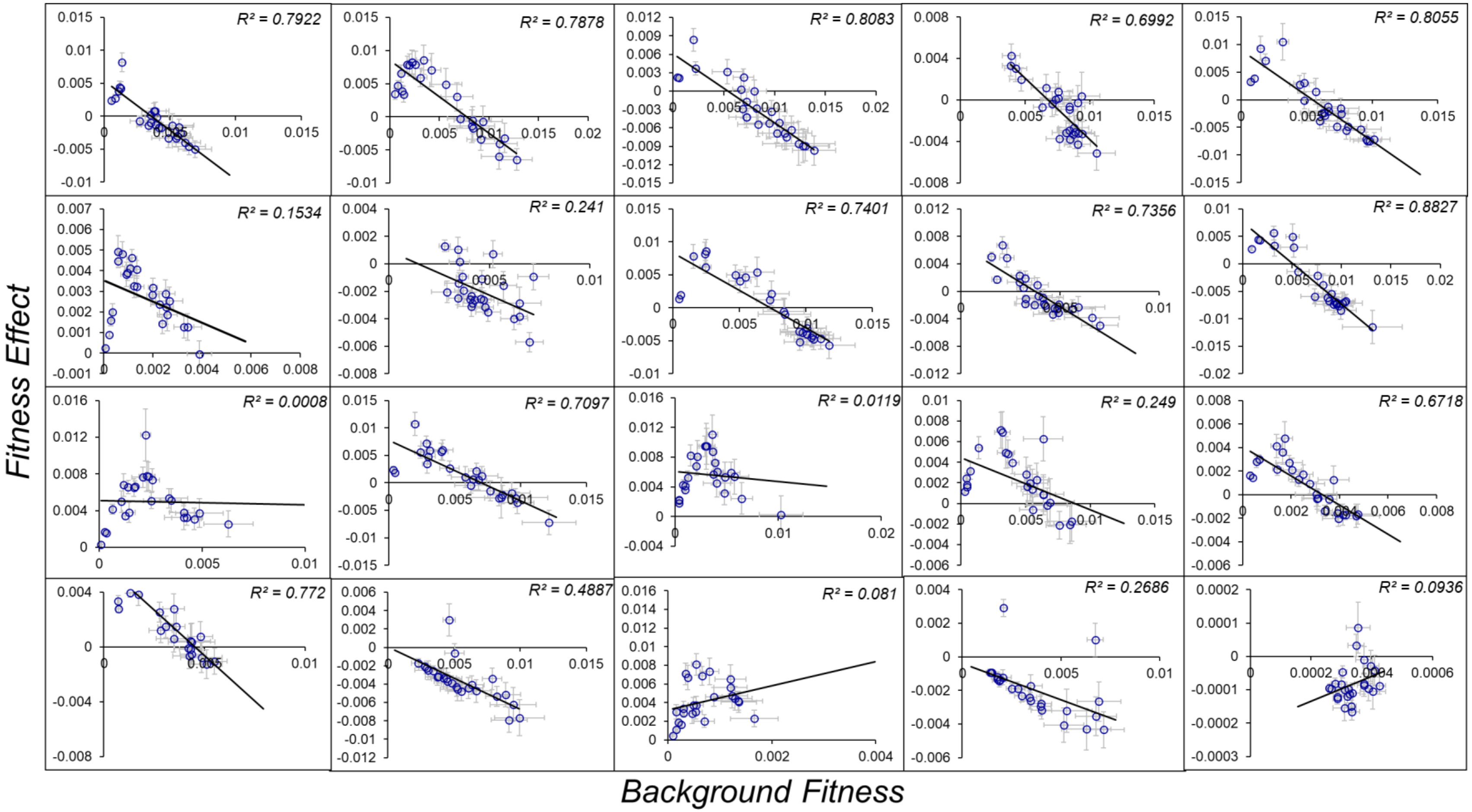

K. Variation in $k_{cat1}$, $k_{cat3}$, and $t_{ON2}$

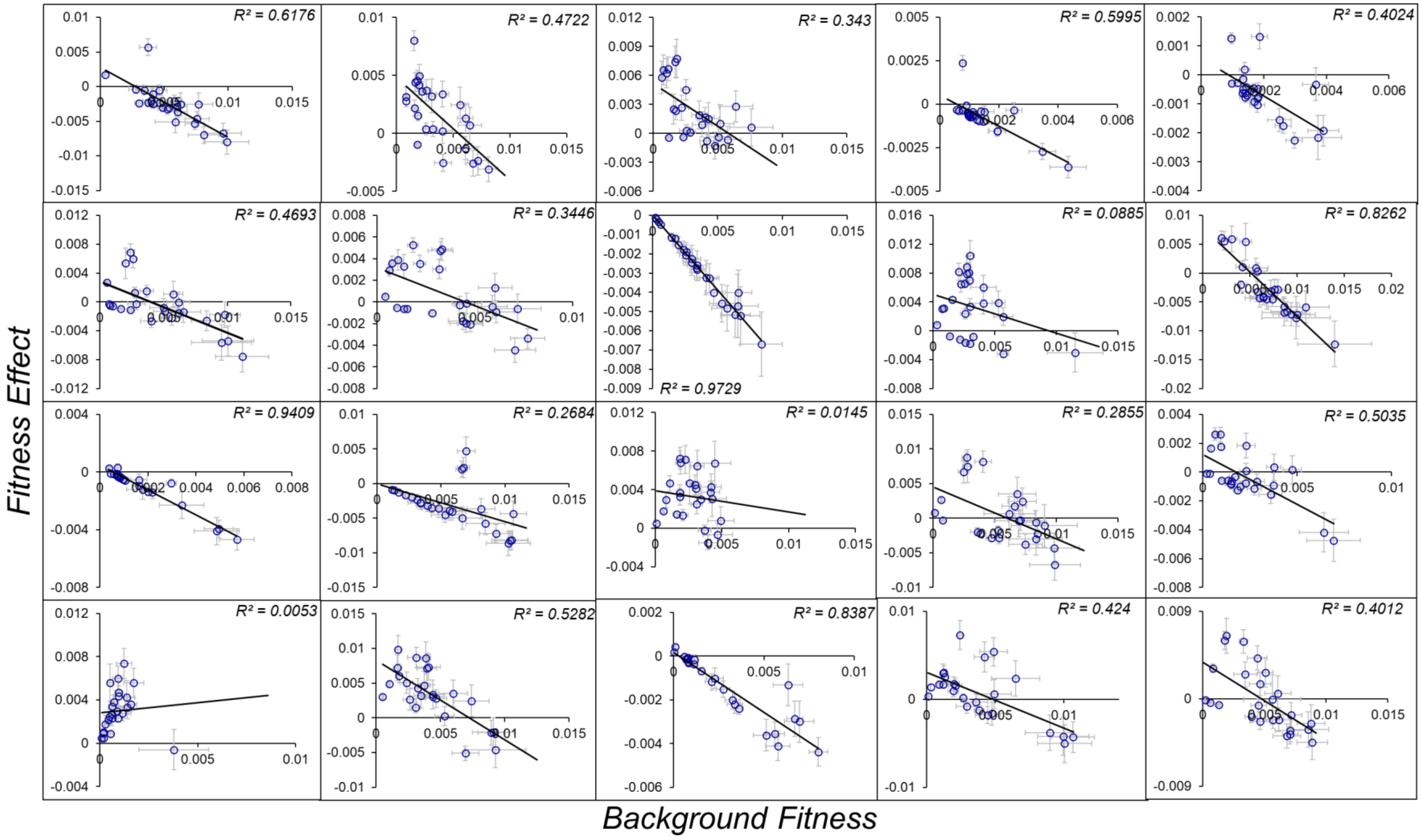

L. Variation in $k_{cat1}$, $t_{ON2}$, and $t_{OFF6}$

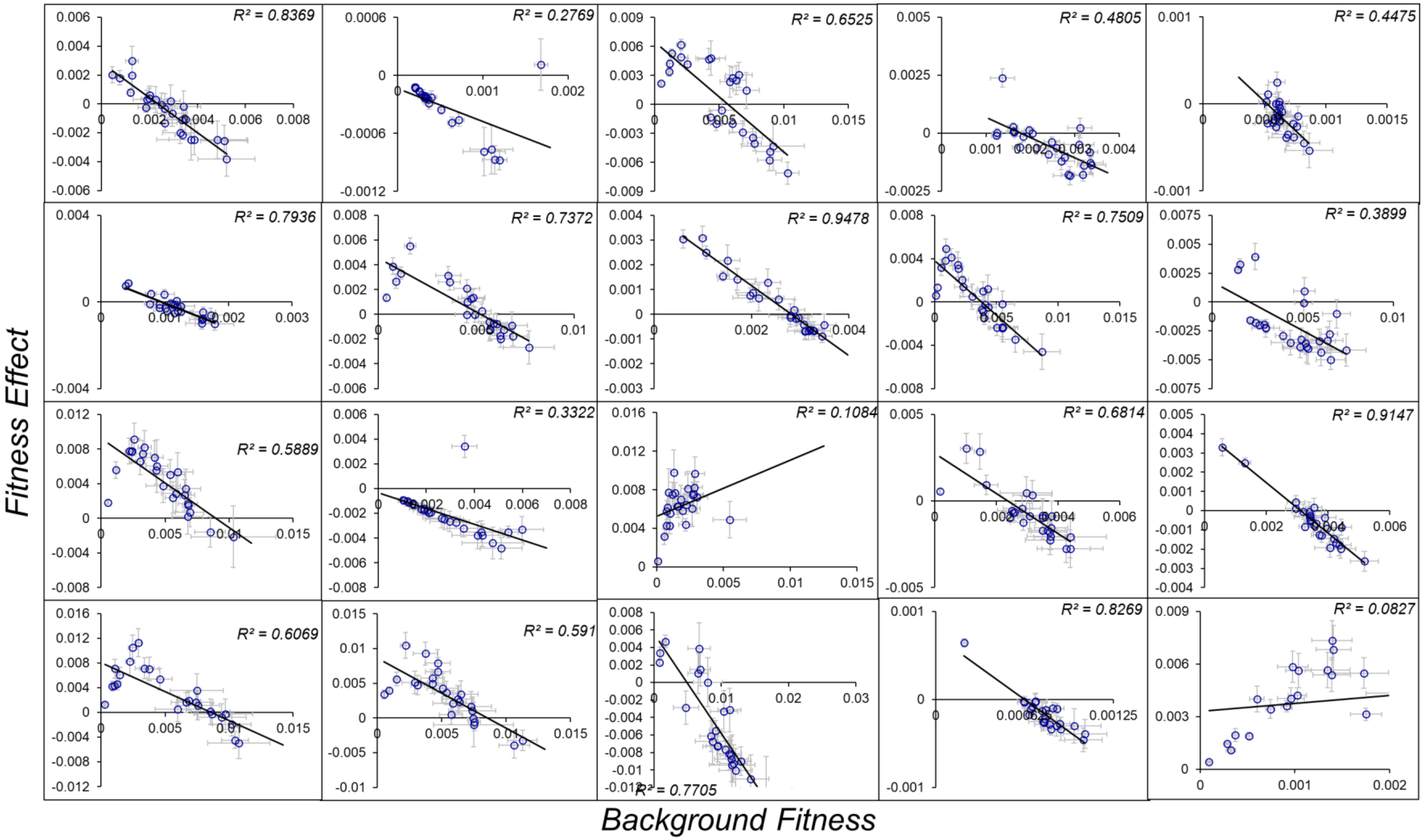

M. Variation in $k_{cat3}$, $t_{ON2}$, and $t_{OFF6}$

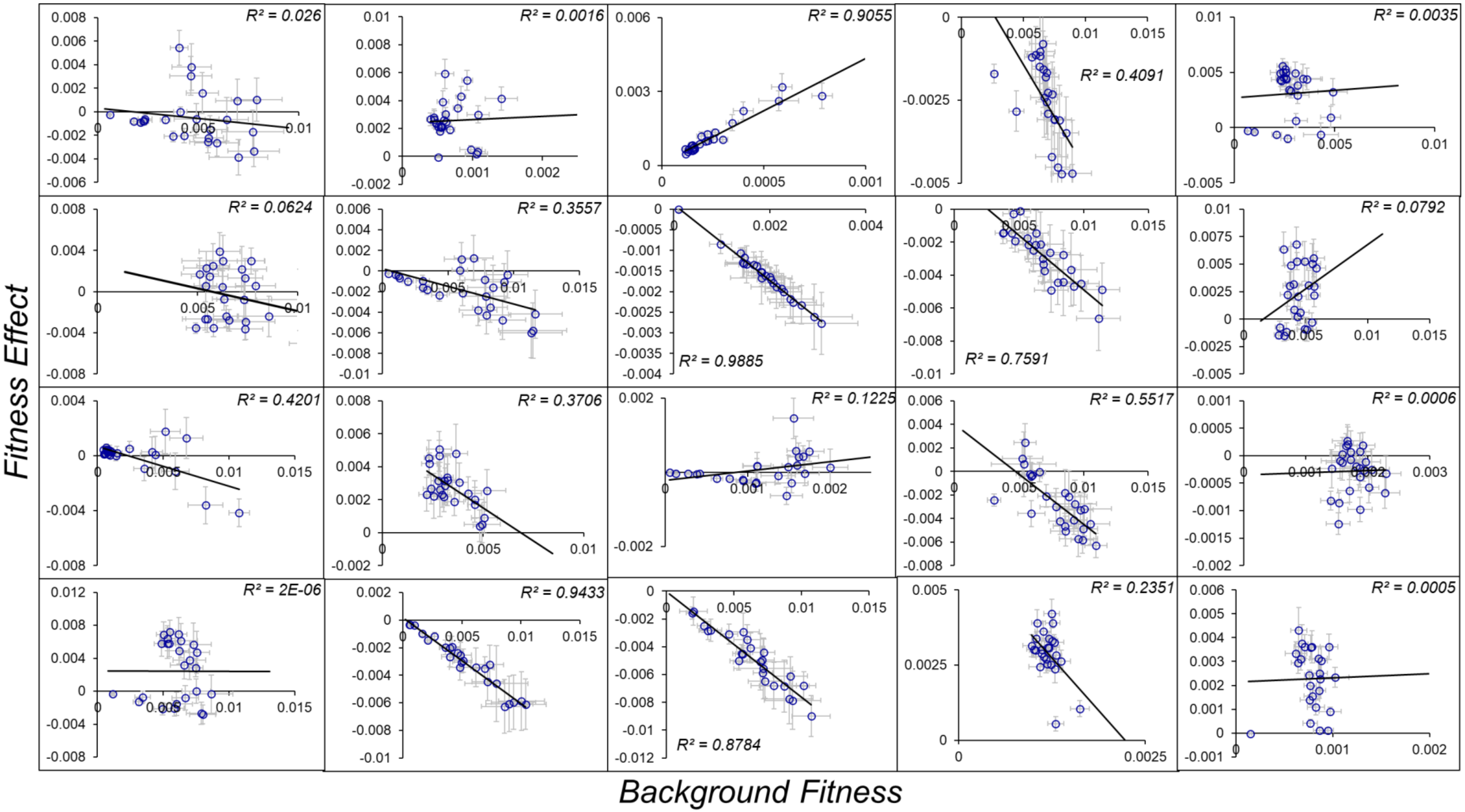

N. Variation in $k_{cat1}$, $k_{cat3}$, and $t_{OFF6}$

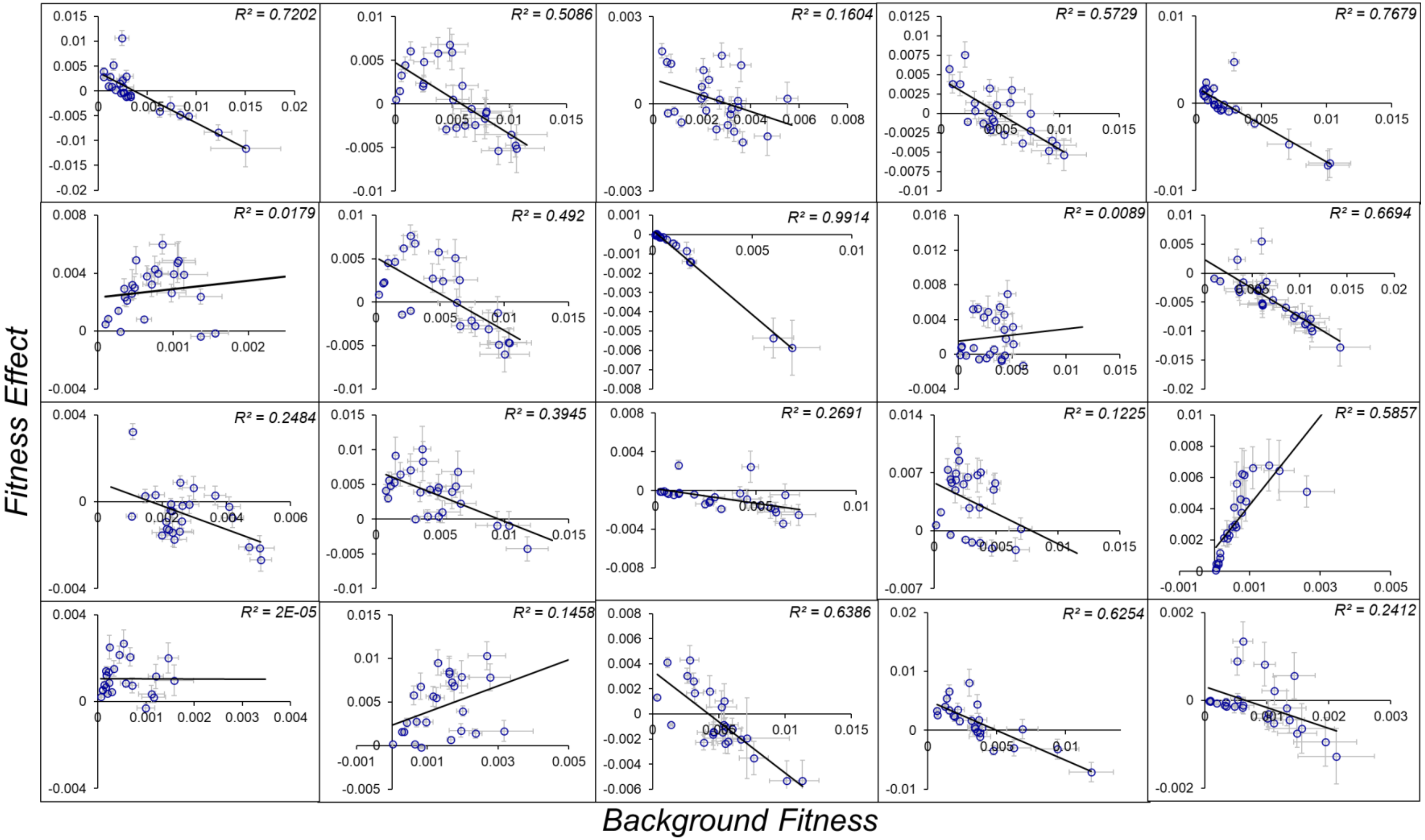

O. Variation in $k_{cat1}$, $k_{cat3}$, $t_{ON2}$, and $t_{OFF6}$

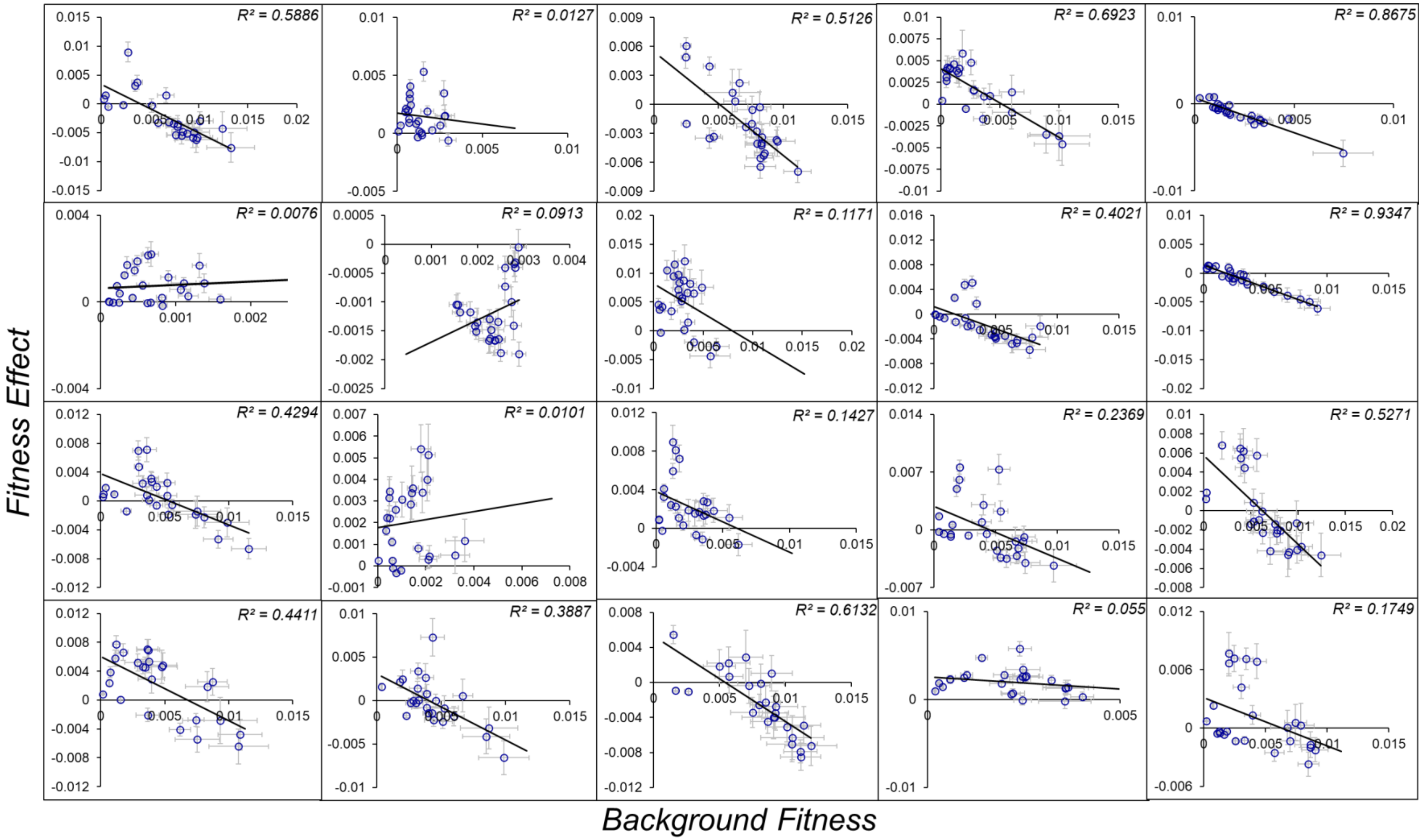

**Figure S8. The fitness effects of mutations can be predicted to different extent using global epistasis approach.** For each type of background variation, we generate twenty *strains*, and twenty-five cells in each *strain* (See Methods). **(A)** to **(D)** indicate the four possibilities where there is one background site variation in $k_{cat1}$ **(A)**, $t_{ON2}$ **(B)**, $k_{cat3}$ **(C)**, and $t_{OFF6}$ **(D)**. Two site variation is represented in **(E)** $k_{cat1}$ and $t_{ON2}$, **(F)** $k_{cat1}$ and $k_{cat3}$, **(G)** $k_{cat1}$ and $t_{OFF6}$, **(H)** $k_{cat3}$ and $t_{ON2}$, **(I)** $t_{ON2}$ and $t_{OFF6}$, and **(J)** $t_{OFF6}$ and $k_{cat3}$. Three site variation is represented in **(K)** $k_{cat1}$, $k_{cat3}$, and $t_{ON2}$, **(L)** $k_{cat1}$, $t_{ON2}$, and $t_{OFF6}$, **(M)** $k_{cat3}$, $t_{ON2}$, and $t_{OFF6}$, and **(N)** $k_{cat1}$, $k_{cat3}$, and $t_{OFF6}$. **(O)** Four site variation is presented with variation in all four parameters $k_{cat1}$, $k_{cat3}$, $t_{ON2}$, and $t_{OFF6}$.

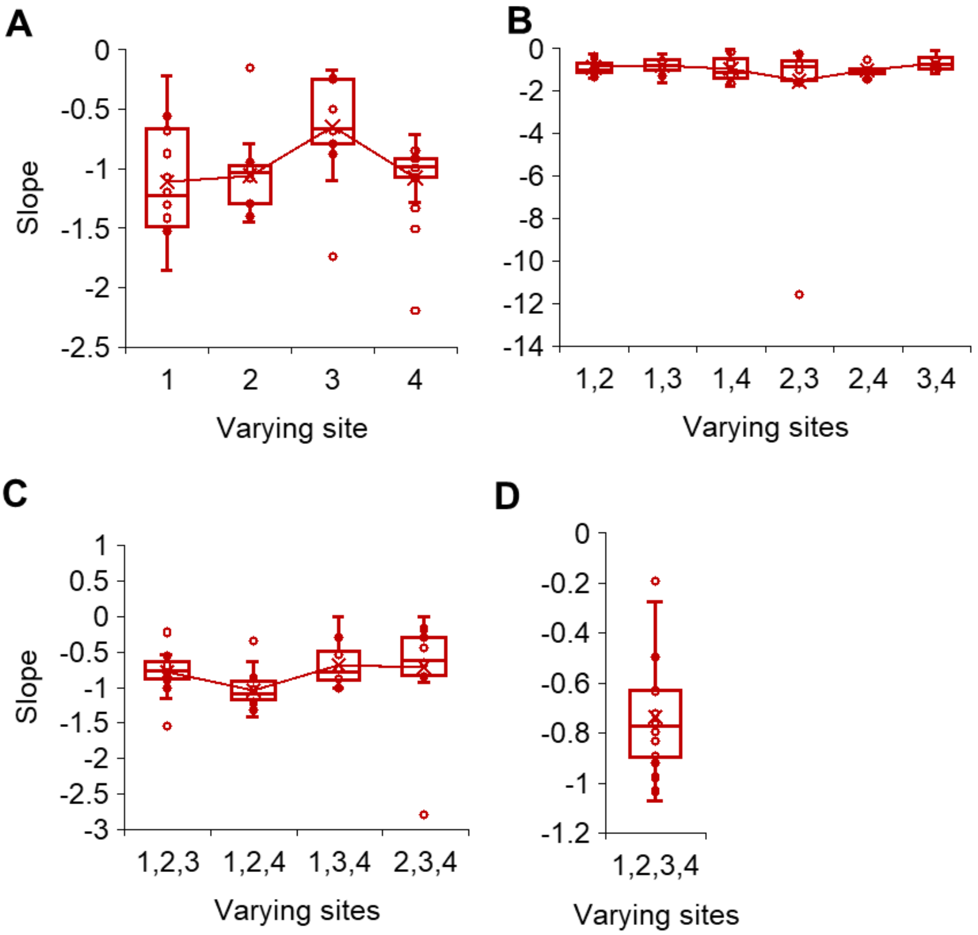


**Figure S9. Distributions of the slopes of negative epistasis.** We quantified the slopes of negative epistasis by performing linear regressions between the fitness effects of a fixed mutation ($0.5 > t_{ON1} > 1$) and the background fitness across different levels of genetic divergence. Specifically, we considered mutation effects in groups of cells that differed from each other at one, two, three, or four background loci, as illustrated in panels (A), (B), (C), and (D), respectively. Parameters varying in the background are shown in the x-axis - $k_{cat1}$, $t_{ON2}$, $k_{cat3}$ and $t_{OFF6}$ are depicted coded as *1,2,3,* and *4*, respectively. Statistical tests were performed to compare the average slopes; the p-values obtained are reported in **Table S2**.

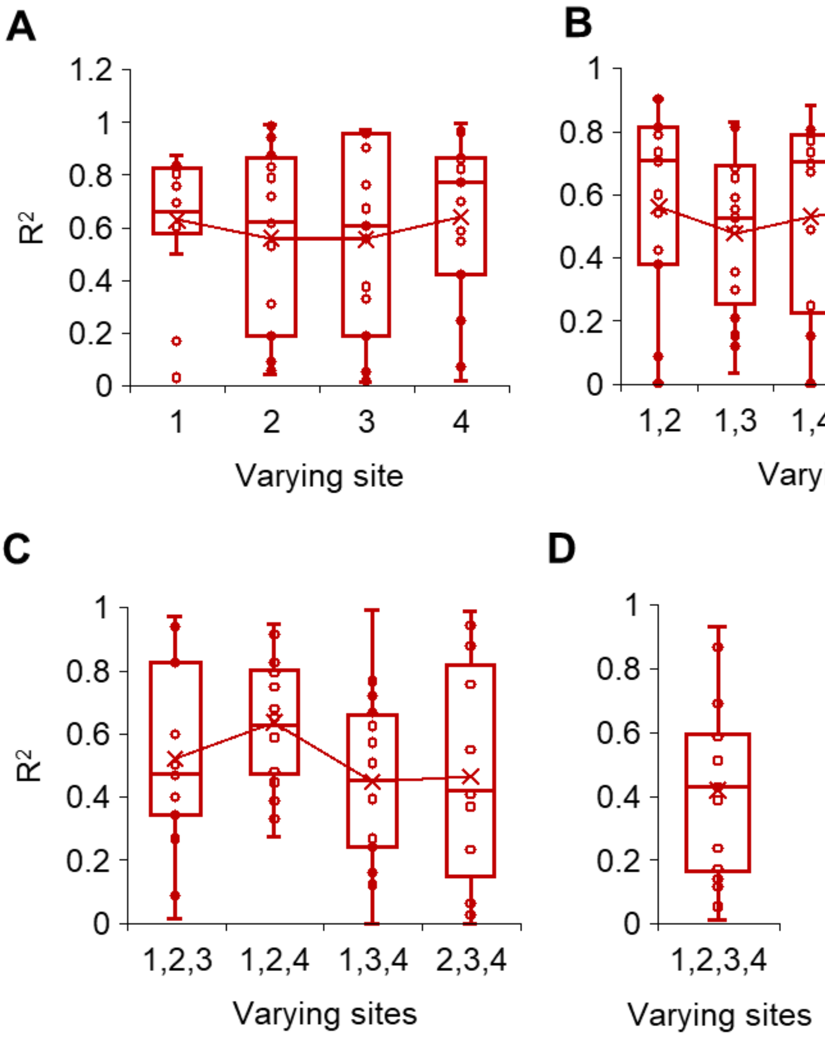


**Figure S10. Distributions of the globality of negative epistasis.** We quantified the $R^2$ of negative epistasis by performing linear regressions between the fitness effects of a fixed mutation ($0.5 > t_{ON1} > 1$) and the background fitness across different levels of genetic divergence. Specifically, we considered mutation effects in groups of cells that differed from each other at one, two, three, or four background loci, as illustrated in panels **(A)**, **(B)**, **(C)**, and **(D)**, respectively. Parameters varying in the background are shown in the x-axis - $k_{cat1}$, $t_{ON2}$, $k_{cat3}$ and $t_{OFF6}$ are depicted coded as *1,2,3,* and *4*, respectively. Statistical tests were performed to compare the average slopes; the p-values obtained are reported in **Table S3**.

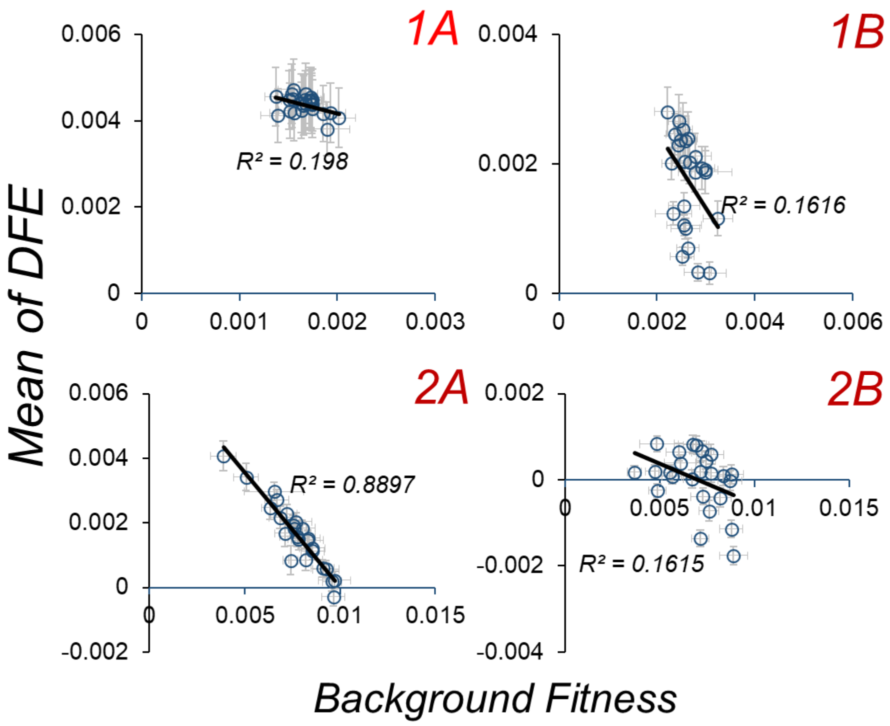


**Figure S11. Strong global epistasis helps predict DFEs.** Using cells with predefined metabolic fluxes, we ask how effective a predictor of the distribution of fitness effects (DFEs) of mutations is background fitness. As shown in above, the mean of the DFE is best predicted in the 2A case, i.e., when mutations occur in the slow second step reaction. In combination with the results shown in Figure 12, it is clear that strong global epistasis makes DFEs predictable.

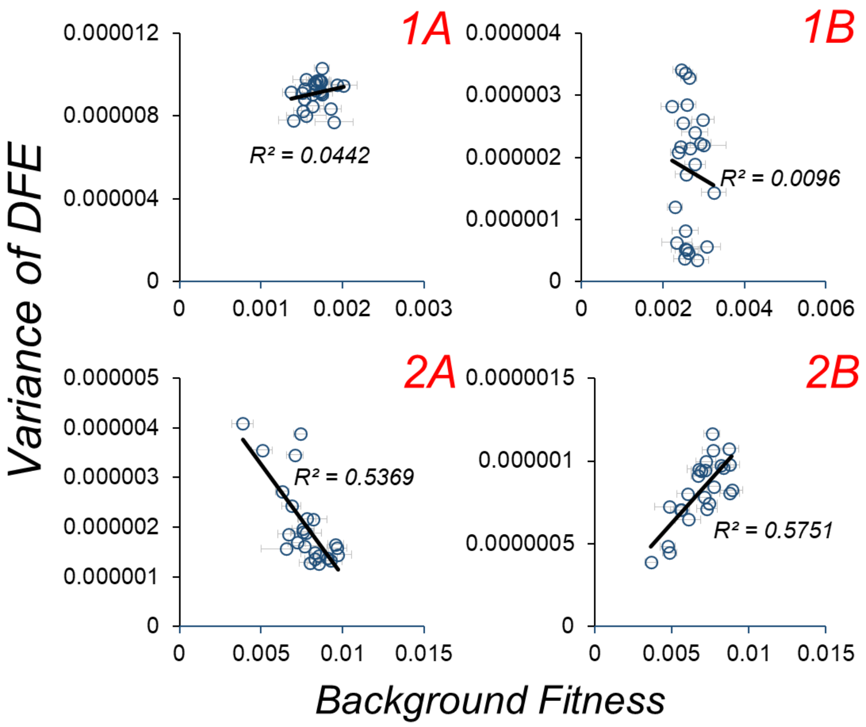


**Figure S12. Metabolic flux modulates the predictability of DFEs.** Using toy cells with predefined metabolic speeds as described in the Methods, we ask if the distribution of fitness effects of mutations is predictable using background fitness. As shown in Figure 13, background fitness serves as an effective predictor of the mean of DFEs in 2A case. However, the variance of the DFE is almost unpredictable for mutations in the first step reactions (both 1A and 1B cases), but is fairly well predictable for mutations in the second step reactions (2A and 2B). Interestingly, changes in variance are not consistent with increase in background fitness, unlike the mean of the DFE, which always decreases with an increase in background fitness.

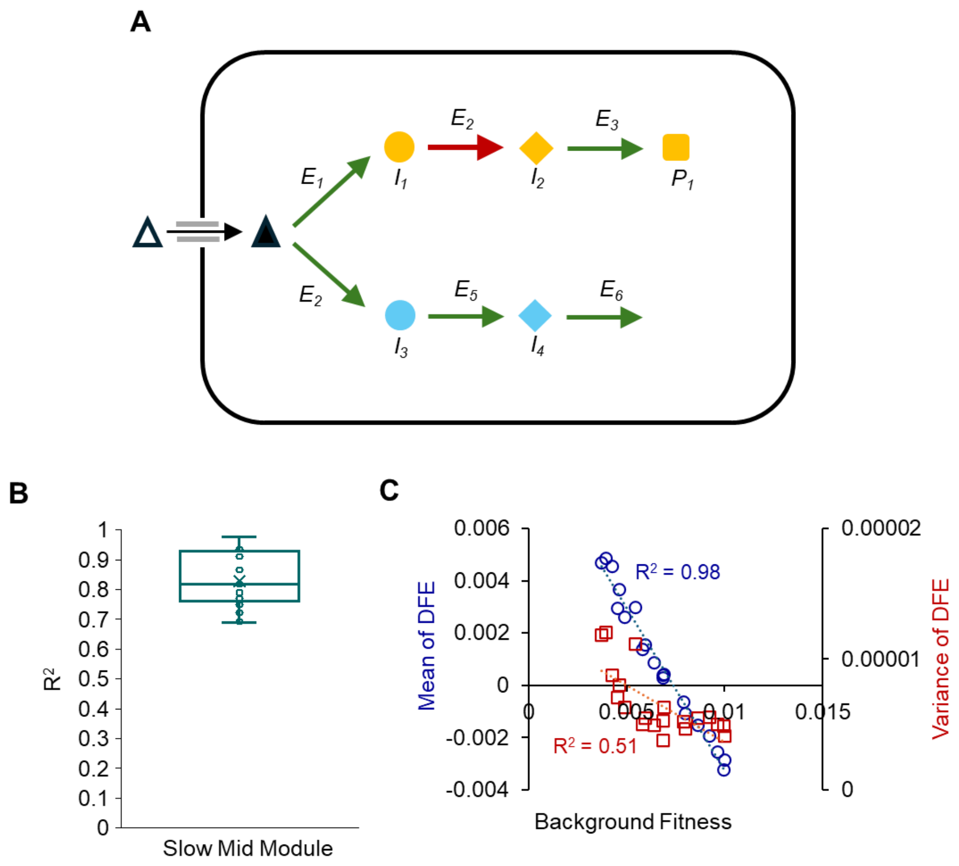


**Figure S13. Mutations in standalone reaction modules exhibit globally epistatic fitness effects. (A)** In the 2×3 architecture, two standalone reaction modules contribute to fitness: one producing the second intermediate ($I_2$ and $I_4$), and the other, the terminal modules producing the threshold metabolite. While **fig. S11** and **S13** show that mutations in the terminal standalone modules exhibit global epistasis, we asked whether non-terminal standalone modules also do so when they are the rate-limiting steps (indicated by red arrow; all other steps were set to be equally fast). **(B)** As shown, the fitness effects of mutations in these non-terminal standalone modules exhibit strong global epistasis, with linear regressions between fitness effect and background fitness yielding an average $R^2$ value of 0.8. **(C)** We further tested whether the distribution of fitness effects (DFE) of these mutations is predictable from background fitness. The mean fitness effect of mutations is well-predicted by background fitness, and background fitness explains ~50% of the variation in the DFE variance.

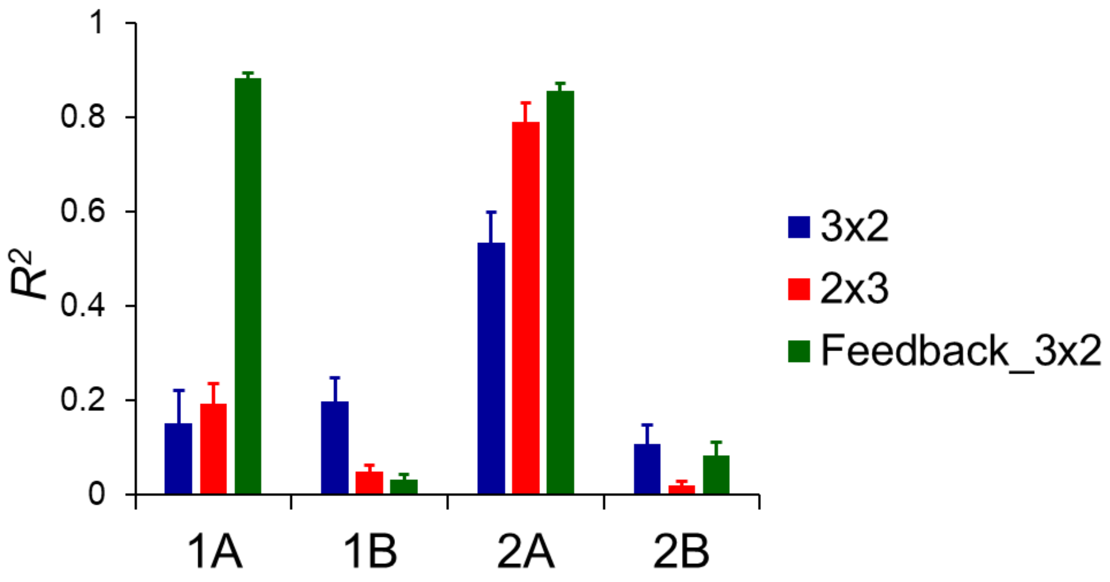


**Figure S14. Globality of epistasis depends on reaction speeds and metabolic control.** Using toy cells with predefined reaction module speeds (as described in the Methods), we investigate if the findings reported in **Figure 3A** holds under different metabolic architectures: a 2×3 configuration (two reaction arms, each with three enzymatic steps), and a 3×2 configuration with strict feedback, where a reaction arm shuts down once its terminal metabolite reaches a threshold concentration. In both scenarios, as shown in the figure, mutations in slow, standalone reaction modules consistently exhibit strong global epistasis, highlighting that the predictability of mutation effects is preserved across architectural variations when metabolic control is localized.

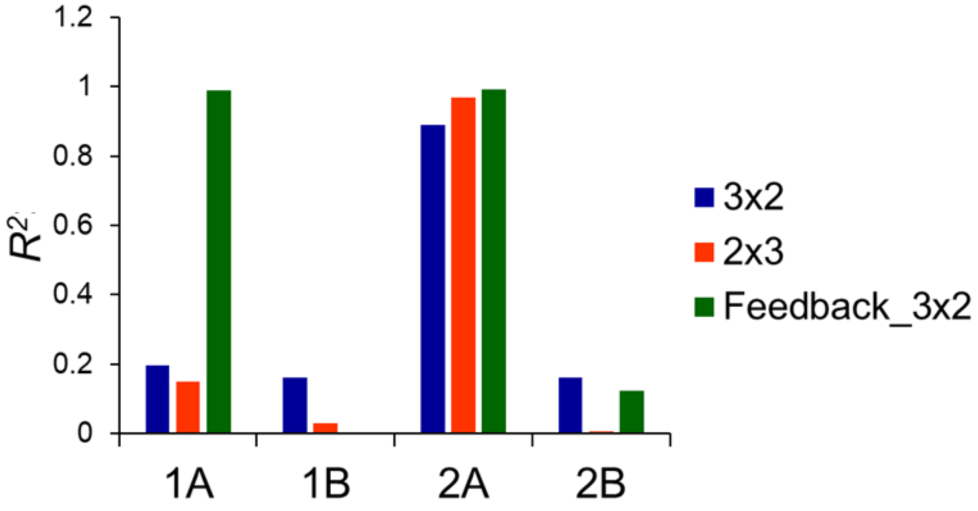


**Figure S15. DFEs and global epistasis.** Using toy cells with predefined reaction module speeds (as described in the Methods), we investigate if the findings reported in **Figure 3F** holds under different metabolic architectures: a 2×3 configuration (two reaction arms, each with three enzymatic steps), and a 3×2 configuration with strict feedback, where a reaction arm shuts down once its terminal metabolite reaches a threshold concentration. In both scenarios, as shown in the figure, mean of the DFEs of mutations in slow, standalone reaction modules are well-predicted using background fitness as indicated by the $R^2$ values.

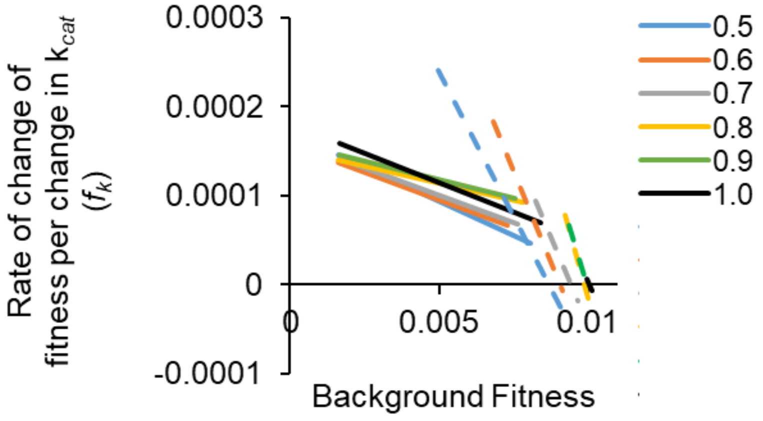


**Figure S16. The rate of change of fitness due to a mutation decreases linearly with the background fitness.** We introduce 4 pre-defined mutation in $k_{cat}$ of the slow reaction modules in six cells each of the 1A and 2A types. As shown by the regression lines in the figure (solid lines – mutations in 1A; dashed lines – mutations in 2A; legends indicate the $t_{ON}$ values in the six different backgrounds), the sensitivity of fitness to changes in $k_{cat}$ correlated well with the fitness of the background in which the mutation occurred ($r_{mean}$ = -0.878 and -0.94 for 1A and 2A, respectively). The slopes of these regressions, however, were more negative for 2A, than 1A, as reported in **Figure 4**.

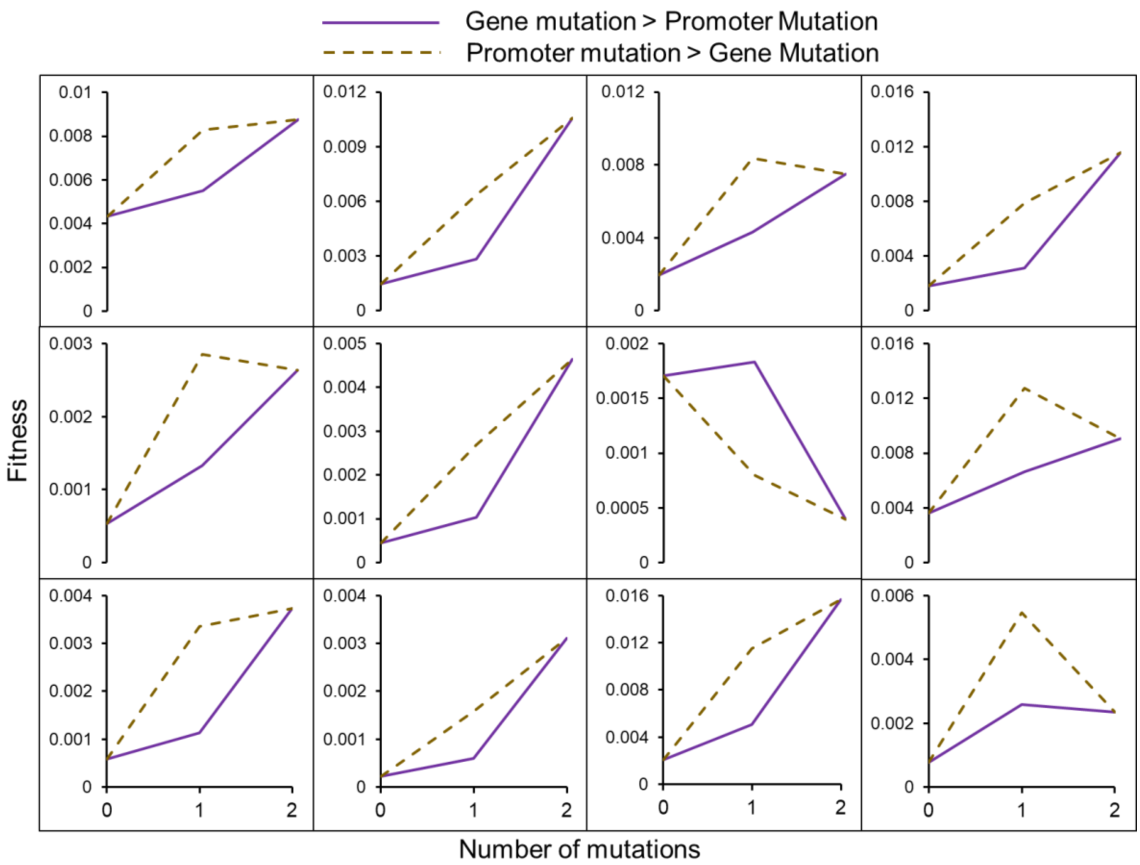


**Figure S17. Promoter-gene landscapes are rugged.** We study the effects of two mutations in a module: a promoter mutation that changes $t_{ON}$ from 0.5 to 1, and gene mutation that changes $k_{cat}$ from 10 to 20. In different genetic backgrounds, we study the fitness changes that result from the two different ways in which these two mutations can be accrued – gene mutation followed by the promoter mutation, and promoter mutation followed by the gene mutation. As shown, gene mutation followed by promoter mutation seems to be a less likely adaptive path, compared to the path where promoter mutation is followed by gene mutation. These results align with experimental protein landscape findings(*74*).

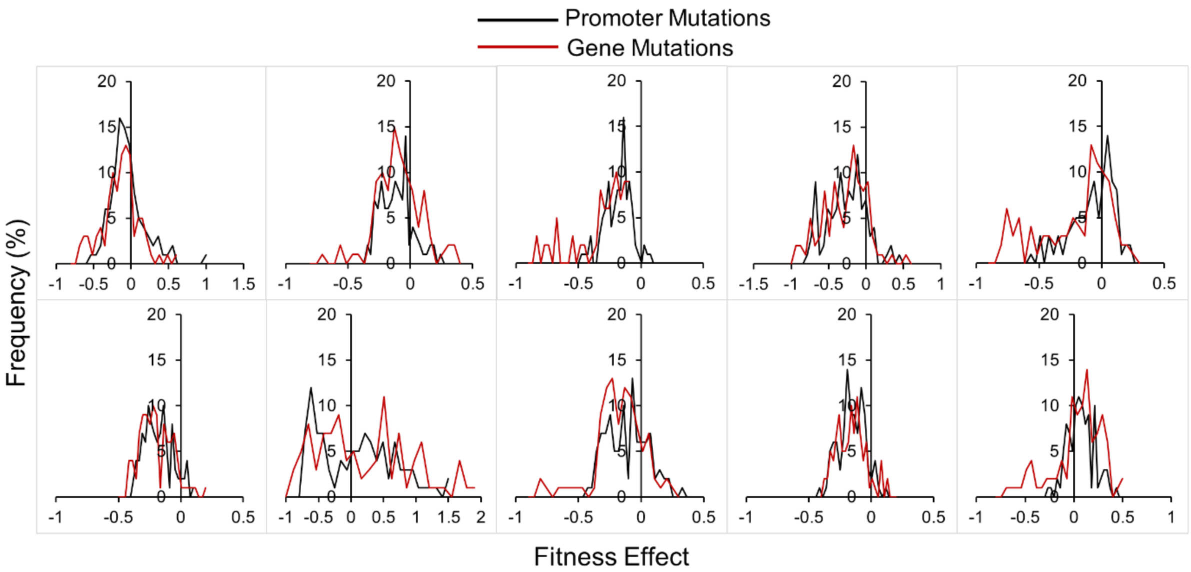


**Figure S18. A module can be altered via mutations in the non-coding or coding region.** In the cells that we model, there are twenty-four parameters which control fitness. We test the effects of a hundred promoter and gene mutations of a module, in ten different cells (different in twenty-two background genetic parameters). In all these genetic backgrounds, we see that the distribution of fitness effects of mutations in the promoter of a module is statistically like that of mutations in the gene (For every pairwise comparison of means using two-tailed t-test, $p$>0.05).

### 3. Supplementary Tables

**Table S1. List of parameters and approximations used in this study.**

| Parameter | Value |
|---|---|
| Threshold metabolites | 1.00E+08 |
| $k_{cat}$ ($s^{-1}$) | U (1,100) |
| $K_M$ (molecules, for a cell of 1$\mu m^3$ volume) | U (6022,602200) |
| $t_{ON}$ (min) | U (0.5,1) |
| $t_{OFF}$ (min) | U (0.5, 5) |
| Mean mRNA lifetime (min) | Exp (25) |
| Mean enzyme lifetime (hours) | Exp (72) |
| Enzyme length | 343 aa |
| Transcription rate | 10nt/s |
| Translation rate | 10aa/s |

**Table S2. Comparisons of the average slopes of regression.** By performing a linear regression of the fitness effects of a mutation (0.5 > $t_{ON1}$ > 1) with the genetic background, we obtain the slopes of regression. Figure S9 shows the distribution of the negative slopes in different scenarios of background variation. This table shows the *p-values* obtained by comparing every pair of mean slopes (using 2-tailed t-test for unequal variance), among the four categories of extents of variation, and among the types that exist within each of the three (one-site, two-site, three-site variation) categories.

**Among extents of variation**

| | |
|---|---|
| 1site-2site | 0.995415 |
| 1site-3site | 0.018271 |
| 1site-4site | 0.001428 |
| 2site-3site | 0.182045 |
| 3site-4site | 0.278506 |
| 2site-4site | 0.051686 |

**Among the one-site variation cases**

| | |
|---|---|
| 1--2 | 0.703062 |
| 1--3 | 0.008558 |
| 1--4 | 0.8176 |
| 2--3 | 0.003055 |
| 2--4 | 0.842075 |
| 3--4 | 0.002917 |

**Among the two-site variation cases**

| | 1,2 | 1,3 | 1,4 | 2,3 | 2,4 | 3,4 |
|---|---|---|---|---|---|---|
| 1,2 | | | | | | |
| 1,3 | 0.571605 | | | | | |
| 1,4 | 0.649185 | 0.357118 | | | | |
| 2,3 | 0.32922 | 0.278917 | 0.389788 | | | |
| 2,4 | 0.217401 | 0.056003 | 0.631994 | 0.443929 | | |
| 3,4 | 0.171133 | 0.390413 | 0.11801 | 0.215921 | 0.007427 | |

**Among the three-site variation cases**

| | 1,2,3 | 1,2,4 | 2,3,4 | 1,3,4 |
|---|---|---|---|---|
| 1,2,3 | | | | |
| 1,2,4 | 0.006306 | | | |
| 2,3,4 | 0.712226 | 0.121751 | | |
| 1,3,4 | 0.281506 | 0.000734 | 0.881139 | |

**Table S3. Comparisons of the average globalities ($R^2$) of the mutation of interest in different types of genetic backgrounds.** By performing a linear regression of the fitness effects of a mutation ($0.5 > t_{ON1} > 1$) with the genetic background, we obtain the globality of epistasis ($R^2$). Figure S10 shows the distribution of the globality in different scenarios of background variation. This table shows the *p-values* obtained by comparing every pair of mean globalities (using 2-tailed t-test for unequal variance), among the four categories of extents of variation, and among the types that exist within each of the three (one-site, two-site, three-site variation) categories.

**Among extents of variation**

| | |
|---|---|
| 1site-2site | 0.25287701 |
| 1site-3site | 0.172650541 |
| 1site-4site | 0.022878314 |
| 2site-3site | 0.775943971 |
| 3site-4site | 0.151356017 |
| 2site-4site | 0.103152992 |

**Among the one-site variation cases**

| | |
|---|---|
| 1--2 | 0.51283882 |
| 1--3 | 0.53941059 |
| 1--4 | 0.90824823 |
| 2--3 | 0.97317517 |
| 2--4 | 0.46147586 |
| 3--4 | 0.48993809 |

**Among the two-site variation cases**

| | 1,2 | 1,3 | 1,4 | 2,3 | 2,4 | 3,4 |
|---|---|---|---|---|---|---|
| 1,2 | | | | | | |
| 1,3 | 0.3664767 | | | | | |
| 1,4 | 0.767366 | 0.572845 | | | | |
| 2,3 | 0.95123512 | 0.399797 | 0.74789 | | | |
| 2,4 | 0.61558767 | 0.16985 | 0.440599 | 0.697627 | | |
| 3,4 | 0.32049883 | 0.77119 | 0.46929 | 0.339715 | 0.166923 | |

**Among the three-site variation cases**

| | 1,2,3 | 1,2,4 | 2,3,4 | 1,3,4 |
|---|---|---|---|---|
| 1,2,3 | | | | |
| 1,2,4 | 0.15697794 | | | |
| 2,3,4 | 0.62659031 | 0.123139 | | |
| 1,3,4 | 0.43678016 | 0.031858 | 0.880845 | |
